# How to evaluate phase differences between trial groups in ongoing electrophysiological signals

**DOI:** 10.1101/061283

**Authors:** Rufin VanRullen

## Abstract

A growing number of studies endeavor to reveal periodicities in sensory and cognitive functions, by comparing the distribution of ongoing (pre-stimulus) oscillatory phases between two (or more) trial groups reflecting distinct experimental outcomes. A systematic relation between the phase of spontaneous electrophysiological signals, before a stimulus is even presented, and the eventual result of sensory or cognitive processing for that stimulus, would be indicative of an intrinsic periodicity in the underlying neural process. Prior studies of phase-dependent perception have used a variety of analytical methods to measure and evaluate phase differences, and there is currently no established standard practice in this field. The present report intends to remediate this need, by systematically comparing the statistical power of various measures of “phase opposition” between two trial groups, in a number of real and simulated experimental situations. Seven measures were evaluated: one parametric test (circular Watson-Williams test), and three distinct measures of phase opposition (phase bifurcation index, phase opposition sum and phase opposition product) combined with two procedures for non-parametric statistical testing (permutation, or a combination of z-score and permutation). While these are obviously not the only existing or conceivable measures, they have all been used in recent studies. All tested methods performed adequately on a previously published dataset (Busch, Dubois & VanRullen, 2009). On a variety of artificially constructed datasets, no single measure was found to surpass all others, but instead the suitability of each measure was contingent on several experimental factors: the time, frequency and depth of oscillatory phase modulation; the absolute and relative amplitudes of post-stimulus event-related potentials for the two trial groups; the absolute and relative trial numbers for the two groups; and the number of permutations used for non-parametric testing. The concurrent use of two phase opposition measures, the parametric Watson-Williams test and a non-parametric test based on summing inter-trial coherence values for the two trial groups, appears to provide the most satisfactory outcome in all situations tested. Matlab code is provided to automatically compute these phase opposition measures.

## 1. Introduction

Science has long sought to determine whether mental processes unfold continuously–like the flow of a river–or discretely over time–like the successive frames of a movie sequence (Stroud, 1956; VanRullen & Koch, 2003). The existence of oscillatory brain rhythms at different spatial and temporal scales, and their demonstrated involvement in numerous sensory and cognitive functions (Buzsaki, 2006), could indeed imply that certain mental processes operate rhythmically, rather than strictly continuously. One convincing way to demonstrate such a rhythmic operation is by showing that the result of a given neural process varies, depending on the exact rhythmic phase at which this process is engaged. Although this procedure has a history dating at least half a century (Callaway & Yeager, 1960; Dustman & Beck, 1965), in recent years there has been a surge of reports of such phase-dependent perception (VanRullen, Busch, Drewes, & Dubois, 2011). The phase of brain oscillations at various frequencies from 2 to 20Hz has been related to trial-by-trial fluctuations in threshold-level perception in the visual (Nunn & Osselton, 1974; Busch, Dubois, & VanRullen, 2009; Mathewson, Gratton, Fabiani, Beck, & Ro, 2009; Busch & VanRullen, 2010; Dugue, Marque, & VanRullen, 2011; Fiebelkorn et al., 2013; Hanslmayr, Volberg, Wimber, Dalal, & Greenlee, 2013), auditory (Rice & Hagstrom, 1989; Ng, Schroeder, & Kayser, 2012; Strauss, Henry, Scharinger, & Obleser, 2015) and somatosensory domains (Ai & Ro, 2014); in supra-threshold perception as measured by reaction times (Callaway & Yeager, 1960; Dustman & Beck, 1965; Drewes & VanRullen, 2011), in oculomotor functions such as saccadic execution (Drewes & VanRullen, 2011; Hamm, Dyckman, McDowell, & Clementz, 2012) and saccadic remapping (McLelland, Lavergne, & VanRullen, 2014), in attention and visual search (Buschman & Miller, 2009; Busch & VanRullen, 2010; Dugue, Marque, & VanRullen, 2015; Landau, Schreyer, van Pelt, & Fries, 2015; Voloh, Valiante, Everling, & Womelsdorf, 2015), in temporal parsing of visual (Varela, Toro, John, & Schwartz, 1981; Chakravarthi & VanRullen, 2012; Cravo, Santos, Reyes, Caetano, & Claessens, 2015; Inyutina, Sun, Wu, & VanRullen, 2015) or somatosensory information (Baumgarten, Schnitzler, & Lange, 2015), in decision-making (Wyart, de Gardelle, Scholl, & Summerfield, 2012), in the top-down influence of predictions and expectations (Arnal, Doelling, & Poeppel, 2015; Han & VanRullen, 2015; Samaha, Bauer, Cimaroli, & Postle, 2015; Ten Oever, van Atteveldt, & Sack, 2015; Sherman, Kanai, Seth, & VanRullen, 2016), in cross-modal integration (van Erp, Philippi, de Winkel, & Werkhoven, 2014) and in short-term memory (Siegel, Warden, & Miller, 2009; Bonnefond & Jensen, 2012; Myers, Stokes, Walther, & Nobre, 2014; Leszczynski, Fell, & Axmacher, 2015). Not surprisingly therefore, large-scale physiological markers of perceptual processing such as ERPs (Dustman & Beck, 1965; Jansen & Brandt, 1991; Haig & Gordon, 1998; Barry et al., 2004; Gruber et al., 2014), stimulus-evoked BOLD responses (Scheeringa, Mazaheri, Bojak, Norris, & Kleinschmidt, 2011) and fMRI network connectivity between areas (Hanslmayr et al., 2013) have also been shown to depend on oscillatory phase at (or just before) the time of stimulus onset.

To quantify the relation between oscillatory phase and a particular cognitive (or physiological) variable, a typical experimental procedure consists in repeating several instances of the same trial, yet leading to different behavioral (or physiological) responses. For example, using a threshold-stimulation procedure, successive presentations of the exact same luminous flash may give rise to a conscious detection of this stimulus in only half of the trials (Busch et al., 2009). If the cognitive function under study (here, visual detection) involves a rhythmic process, then the two trial groups (detected vs. undetected) might be found to differ in the distribution of oscillatory phases at the critical frequency, around the time of stimulus onset. Statistically evaluating this difference in oscillatory phase angle can be (and has been) done in various ways. For example, parametric tests of differences in circular distributions are available (e.g. circular Watson-Williams test), that are equivalent to a t-test for linear data. Such tests are relatively easy to perform (Baumgarten et al., 2015; Samaha et al., 2015), but require the data to verify specific constraints (e.g. normality). It is also possible to construct ad hoc measures of phase opposition between the two trial groups, and evaluate their significance using non-parametric statistics. For this, it is helpful to recognize that, if phase influences the trial outcome, then the inter-trial phase coherence (ITC) of each trial group should exceed the overall inter-trial phase coherence. Thus, phase opposition measures generally involve a combination (sum or product) of ITC for each trial group, appropriately corrected (by subtraction or division) to remove the overall ITC. For example, the ‘phase bifurcation index’ (PBI) introduced by Busch et al (2009), and employed several times since (Hamm et al., 2012; Ng et al., 2012; Auksztulewicz & Blankenburg, 2013; Hanslmayr et al., 2013; Manasseh et al., 2013; Rana, Vaina, & Hamalainen, 2013; Diederich, Schomburg, & van Vugt, 2014; Park, Correia, Ducorps, & Tallon-Baudry, 2014; Li et al., 2015; Shou & Ding, 2015; Strauss et al., 2015; van Diepen, Cohen, Denys, & Mazaheri, 2015; Batterink, Creery, & Paller, 2016), was based on this principle. Other analogous procedures have been described, however (Drewes & VanRullen, 2011; Dugue et al., 2011; VanRullen et al., 2011; Dugue et al., 2015; Han & VanRullen, 2015), and there exists no systematic comparison between these various measures and, consequently, no accepted practice in this field.

The present study aims to compare seven variants of phase opposition measures that have been used in previous published studies. By applying all measures to the same experimental datasets, and by systematically varying key experimental parameters in artificially constructed datasets, we can gain insight about the relative merit of each phase opposition measure, and the conditions under which it should (or should not) be applied. We hasten to note that a number of other measures might already exist and that many more could also be conceived for similar purposes—in other words, this comparison is not intended to be exhaustive, but it should at least contribute to organizing a significant portion of the existing literature.

## 2. Methods

### 2.1 Experimental assumptions

In all following analyses, we shall work on (real or artificial) experimental datasets for which two possible outcomes A and B have been recorded in otherwise identical trials, resulting in two trial groups A and B. (It is worth noting, however, that both parametric and non-parametric measures of phase opposition can readily be extended to situations where the number of possible outcomes is larger than 2.) Each trial is also associated with a time-varying electrophysiological signal, which may represent EEG or MEG from a specific sensor, an intracranial electrode, etc.

We further assume that stimulus onset (t=0) for every trial occurs in a temporally unpredictable manner (e.g. because of randomized inter-trial intervals). This assumption ensures that we consider truly ongoing brain activity: even if certain brain states fluctuate rhythmically, each phase of this fluctuation would be equally likely to be sampled at stimulus onset. In other words, for every frequency, the pre-stimulus phases across trials can be assumed to be sampled from a uniform distribution^1^ Note that this is not necessarily the case concerning the post-stimulus period, as stimulus-evoked activity (event-related potential or ‘ERP’) may affect the observed distribution of phases. Further, the temporal smearing of oscillatory signals caused by window-based time-frequency analysis methods (e.g. wavelet transform) can potentially result in this influence of stimulus-evoked activity being already visible in the pre-stimulus period, a phenomenon that will be explored in various upcoming simulations.

A major assumption for our analyses is that the phase of oscillatory activity at one particular time and frequency point has a significant influence on experimental outcome (i.e., trial assignment to group A or B). Consequently, the purpose of our phase opposition analyses is to reveal the oscillatory frequency involved. That is, we place ourselves in the situation of an experimenter trying to determine whether task outcome is rhythmically modulated by any oscillatory signal, and if yes, at what frequency. Phase opposition must therefore be evaluated for all time and frequency points (‘time-frequency’ analysis). Although we do not explore or simulate ‘control’ datasets in which no phase modulation is present, the likelihood of detecting phase opposition at an incorrect time and/or frequency can be taken as a measure of false alarm rate or baseline (chance-level) performance for our procedures. (Note finally that, in specific cases, experimenters may have strong a priori hypotheses about the exact rhythmic frequency involved; in such cases, which we do not address here, other analyses procedures may be warranted.)

When multiple datasets are recorded for a given experimental situation (e.g. multiple subjects in a given experiment), we assume that the rhythmic modulation that we aim to reveal occurs around the same time and frequency for all datasets. However, the exact phase value favoring outcome A vs. B may or may not be the same across datasets/subjects. (Different reasons may justify this assumption: various conduction delays resulting in shifts of optimal phases; differences in cortical folding resulting in signal polarity reversals, etc. For all these reasons, while the absolute phase of local cortical oscillations at the level of a given neuronal ensemble is obviously important for its operation, we generally consider the absolute phase *recorded at the scalp* to be irrelevant, and focus instead on the *relative* phase between the two trial outcomes). Consequently, our analyses shall evaluate phase opposition between trial groups (‘single-trial’ analysis), independently for each dataset (whose results can later be combined), rather than measuring phase opposition across datasets/subjects (between the mean phase angles of each trial group, i.e., a ‘group-level’ analysis).

### 2.2 Phase opposition measures

Having fixed the experimental conditions common to all analyses, we now introduce the different tests of phase opposition that will be compared. All of these tests are based on a comparison of inter-trial phase Coherence (ITC) measured over all trials (serving as a baseline) vs. ITC measured separately for each trial group (A and B). The null hypothesis tested is, therefore, that the ITC of each trial group exceeds the overall ITC. That is, if ω_i_ is a complex number representing the oscillatory signal (at a given time and frequency) for trial i (with | ω_i_ | and angle(ω_i_), respectively, representing oscillatory amplitude and phase), then ITC (Lachaux, Rodriguez, Martinerie, & Varela, 1999) is defined as:

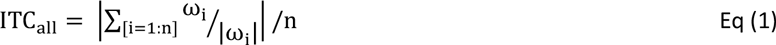

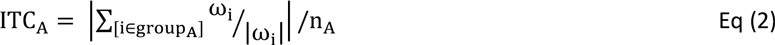

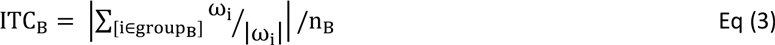

with n_A_ and n_B_ corresponding to the numbers of trials in groups A and B, respectively, and n=n_A_+n_B_.

The various tests simply differ in the ways they combine the above three ITC values, as well as their derivation of statistical significance.

#### 2.2.1 Circular Watson-Williams test

The circular Watson-Williams test is a two-sample test for equal means, equivalent to a two-sample t-test for linear data. It assumes that each set of phases to be compared follows a von Mises circular distribution, and that the two distributions share a common concentration parameter κ. For our purposes, we adapted the implementation provided in the Matlab “circStat” (circular statistics) toolbox (Berens, 2009), in order to process multi-dimensional datasets (time-frequency matrices). This function implements the procedure described by Zar (Zar, 1999), where the test statistic F defined as:

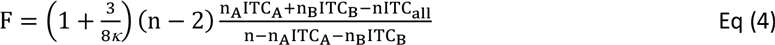

follows a F distribution with (1,n-2) degrees of freedom. Since this parametric test statistic can be directly related to the corresponding p-value (by means of the F cumulative distribution function), no further statistical analysis was required for this test.

#### 2.2.2 Phase bifurcation index (PBI)

We measured the phase bifurcation index as described in Busch et al (2009):

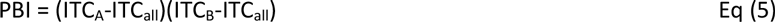

The PBI is bounded between −1 and 1. It takes positive values when the ITC of each trial group exceeds the overall ITC, our main situation of interest. However, PBI can also become negative if the ITC of one trial group happens to fall below the overall ITC (a situation that may occur due to measurement noise, but also due to differences in ERPs shape or amplitude across the trial groups). Finally, PBI can also turn positive in rare cases where (e.g. due to measurement noise) both trial groups’ ITC values are smaller than the overall ITC. It is easy to understand, therefore, that PBI can prove a rather volatile measure. Nonetheless, this volatility should also be present in surrogate distributions of PBI calculated under the null hypothesis (see 2.3. Statistical Analysis), and thus it need not thwart the statistical power of the PBI measure.

#### 2.2.3 Phase opposition product (POP)

A simple modification of the phase bifurcation index can presumably help remove some of its volatility, by subtracting the baseline quantity ITC_all_ outside, rather than inside the product operation. Indeed, in this case a small change in either ITC_A_ or ITC_B_ is less likely to produce a sign reversal of the phase opposition measure. This was the logic employed in a recent study (Han & VanRullen, 2015). That is:

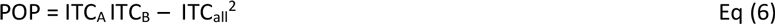

It is worth noting also that the baseline term −ITC_all_^2^ in Eq (6) will be identical in any permutation of the trial assignments (since the overall ITC does not change). Thus, this correction term, which is helpful to display and interpret raw phase opposition measures, can be simply discarded in the statistical analysis when surrogate distributions of POP are calculated under the null hypothesis (see 2.3. Statistical Analysis). This would not be the case for the PBI measure, because the correction in that case is not a mere subtraction.

#### 2.2.4 Phase opposition sum (POS)

Following a number of recent studies (Drewes & VanRullen, 2011; Dugue et al., 2011; McLelland et al., 2014; Bompas, Sumner, Muthumumaraswamy, Singh, & Gilchrist, 2015; Dugue et al., 2015; Inyutina et al., 2015; Sherman et al., 2016), we also computed the simple sum of ITC_A_ and ITC_B_ (again corrected by subtracting the baseline ITC_all_). That is:

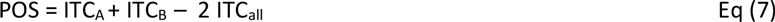

Just like the previous 2 measures, POS will be positive when the ITC of each trial group exceeds the overall ITC. However, it can be argued that using a sum instead of a product renders this measure more stable than the other two. As previously, the subtractive correction term, important for display and interpretation, can be omitted during the permutation procedure (see 2.3. Statistical Analysis).

### 2.3 Statistical analysis

From a given dataset, we computed seven time-frequency maps of p-values, corresponding to seven distinct ways of evaluating phase opposition, as illustrated in Figure 1. The electrophysiological signal recorded on every trial was subjected to a time-frequency transform (‘timefreq’ function from the Matlab EEGlab toolbox, using the ‘wavelet’ option, and frequencies increasing logarithmically from 2 to 50Hz while the number of cycles in each wavelet increases linearly from 2 to 15 cycles). At each time and frequency point, the values ITC_A_, ITC_B_ and ITC_all_ were computed, and from these values, four distinct time-frequency maps were obtained. One map directly contained the p-values resulting from the circular Watson-Williams test (Eq 4); this corresponds to the procedure employed for example by Baumgarten et al (2015). The remaining three maps stored the phase opposition measures PBI, POP and POS (Eqs 5–7). In order to assess the statistical significance of these measures, two non-parametric permutation procedures were employed.

#### 2.3.1 Permutation test

For a given dataset, the trial assignment to group A or B was randomly permuted a number of times (n_perm_; here n_perm_=1,000, except when stated otherwise). The phase opposition measures PBI, POP and POS were recomputed after every permutation. For each of these three measures, at each time-frequency point, the final p-value assigned was the proportion of permutations that yielded a higher measure than in the original dataset, or 1/2n_perm_, whichever was highest. Note that low numbers of permutations limit the range of p-values that can be obtained, which can prove problematic, for example when a correction for multiple comparisons across time and frequency points is needed (e.g., with n_perm_=1000, no result can survive a Bonferroni correction when the number of time-frequency points is larger than 100). On the other hand, permutations of time-frequency data are computationally intensive, and it is thus not always possible to reach adequate values of n_perm_.

#### 2.3.2 Permutation + z-score test

To circumvent this problem, we have suggested a streamlined procedure (Drewes & VanRullen, 2011; Dugue et al., 2011; McLelland et al., 2014; Dugue et al., 2015; Inyutina et al., 2015; Sherman et al., 2016), in which a relatively low number of permutations is used to characterize the mean and standard deviation of the null-hypothesis distribution. The true value from the original dataset is then compared against the null-hypothesis distribution by means of a z-score. In other words, for each of the three phase opposition measures (PBI, POP, POS), for each time-frequency point, the difference between the original dataset and the mean of all permutations was expressed in units of standard deviation (across all permutations). The resulting time-frequency map of p-values was obtained by means of the normal cumulative distribution function.

**Figure 1.**
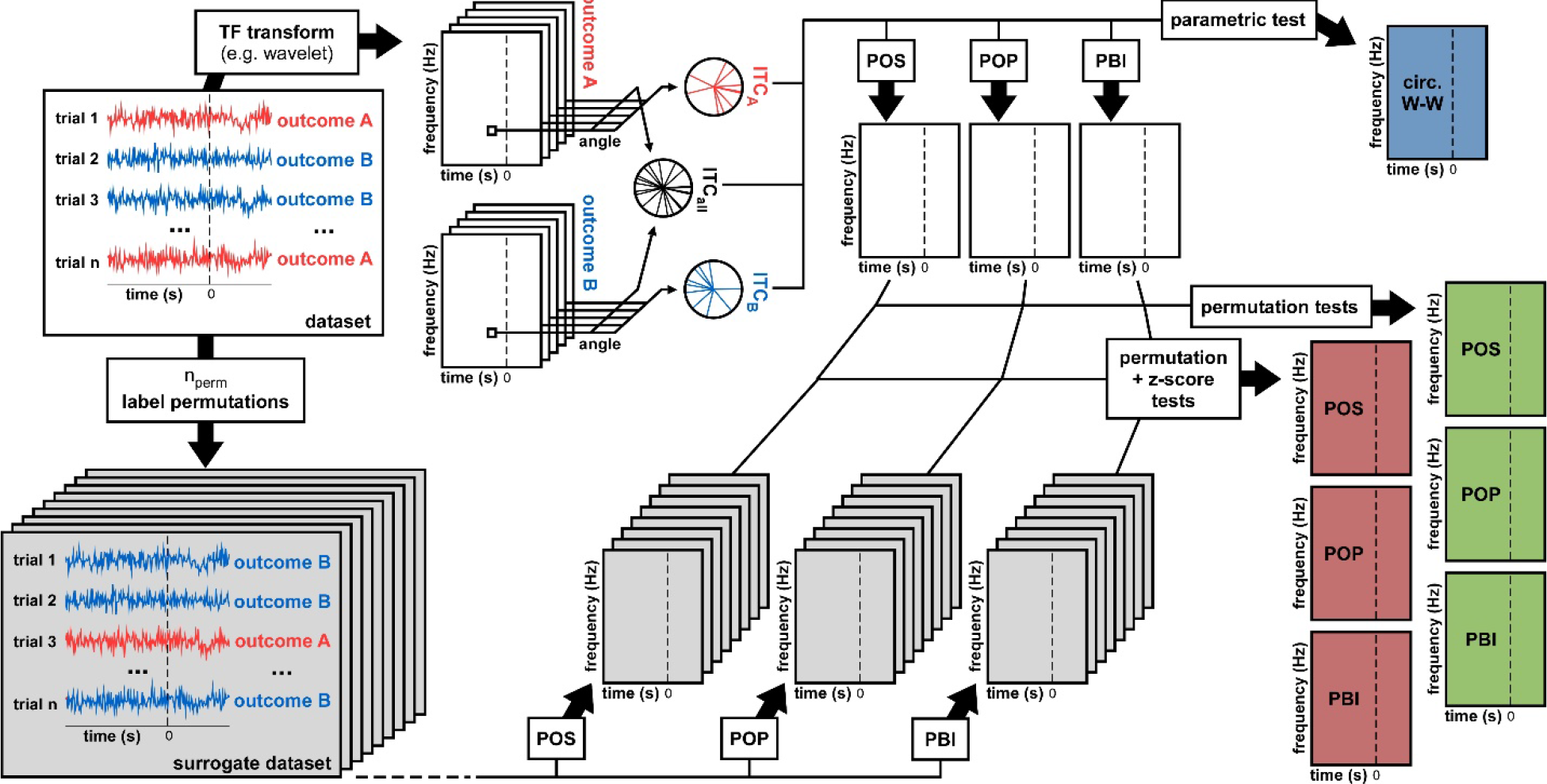
Data analysis methods. From a dataset we derived 7 statistical measures of phase opposition. A time-frequency transform was used to extract oscillatory phase for each trial, time point, and frequency. Inter-trial coherence values were then computed for each trial group (outcome A vs B) as well as for the entire dataset (both outcomes pooled). Using these 3 ITC values, the circular Watson-Williams parametric test directly yielded a time-frequency map of p-values (top-right, blue time-frequency map). The same 3 ITC values were used to calculate time-frequency maps of the phase opposition measures POS, POP and PBI. To determine the statistical significance of these measures, two distinct non-parametric procedures were applied. The first consisted in randomly permuting the trial labels (assignment of outcome A vs. B), and for each permutation recalculating POS, POP and PBI. The ranking (percentile) of the original dataset values against these null-hypothesis distributions could be used as a p-value (right hand-side, green time-frequency maps). Alternately, the null-hypothesis distributions could be summarized by their mean and standard deviation (across permutations), the values of the original dataset could then be expressed as a standardized z-score against the null distribution, and a p-value assigned using the normal cumulative distribution function (right hand-side, red time-frequency maps).

### 2.4 Test performance on real experimental dataset

The above statistical analysis procedure describes how the seven p-value maps of phase opposition are derived from a given dataset (Figure 1). It remains to be seen how each of these seven measures will perform in situations where the “ground truth” is known (or assumed to be known). Is a significant phase opposition detected at the proper time and frequency point? Are there other, spuriously significant phase opposition values detected at erroneous time-frequency points? We first addressed these questions using a previously published experimental dataset (Busch et al., 2009) for which the existence of phase opposition around 7Hz and 120ms pre-stimulus could be assumed as “ground truth” (insofar as previous publication can be considered a mark of reliability).

#### 2.4.1 p-value combination

As the experiment had been performed for multiple observers (N=12), each of whom contributed one dataset for the statistical analysis described in section 2.3 and in Figure 1, it was necessary to combine the results (time-frequency maps of p-values) across observers. For this purpose, we used the method described by Stouffer (Stouffer, Suchman, DeVinney, Star, & Williams, 1949), whereby each p-value is turned into an equivalent z-score (using the inverse normal cumulative distribution function Φ^−1^), the z-scores are combined across observers and finally turned back into probabilities (using the normal cumulative distribution function Φ), that is:

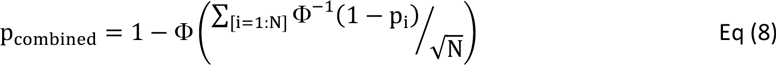

#### 2.4.2 AUC

Given the conclusions of the previously published experiment (Busch et al., 2009), we considered that a phase opposition method was efficient if it detected significant pre-stimulus phase opposition *within* the expected time-frequency region (here, defined as a rectangular region of extent 140ms and 4Hz, centered on the expected phase opposition peak, at 120ms pre-stimulus and 7Hz) but not *outside* of this expected region. Since the answer can strongly depend on the choice of a significance threshold (too conservative and no phase opposition can be detected at all; too liberal and phase opposition materializes at all times and frequencies), we used an ROC procedure (Receiver-Operator Characteristic) to measure efficiency in an unbiased manner. Each possible significance threshold was used alternately, and each time the proportions of ‘hits’ (significant p-values inside the target time-frequency region) and ‘false alarms’ (significant p-values before stimulus onset but outside the target region) were recorded. Plotting hit rate against false alarm rate produces the ROC curve, and the area under this curve (AUC for ‘Area Under the ROC Curve’) can serve as a measure of sensitivity, with chance level at 0.5 and maximal performance at 1. This AUC was calculated for each of the seven phase opposition measures.

### 2.5 Artificial datasets creation

To determine and compare the sensitivity of each phase opposition measure in a variety of well-controlled situations, we also created artificial datasets, for which the parameters of oscillatory modulation could be precisely ascertained (Figure 2). Just like with real experimental data, an artificial dataset was made of n trials, each of which had a (simulated) electrophysiological signal associated with an experimental outcome A or B. The electrophysiological data of each trial was randomly initialized with white noise^2^. This signal was bandpass filtered around a specific frequency f (different for each simulation), and a Hilbert transform converted the resulting oscillatory waveform into complex values. The phase angle was extracted at a critical time point t (different for each simulation), and the trial outcome was decided on the basis of this phase value. The probability of outcome A was a cosine function of this phase angle^3^:

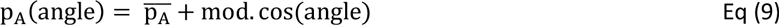

where 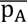 is the overall probability of outcome A, and mod is the depth of the phase modulation (both variable parameters). The probability of outcome B was, therefore, p_B_=1-p_A_.

Once the trial outcome was decided, a noisy ERP-like signal was added to each trial, with (potentially) different amplitudes for trial groups A and B. The shape of the ERP was bimodal, with a first positive peak (a ‘P1’ component) and a second negative one (a ‘N1’ component). The standard P1 was a Gaussian waveform, slightly different on each trial, with peak latency μ drawn from a normal distribution (mean=65ms, standard deviation=10ms) and temporal extent ct drawn from a normal distribution (mean=8.33ms, standard deviation=1.66ms; the waveform was zero-padded outside a window duration of 6σ); the maximal amplitude of the P1 was drawn from a normal distribution (mean=1, standard deviation=0.5, arbitrary units). Similarly, the standard N1 was a negative Gaussian waveform, slightly different on each trial, with peak latency μ drawn from a normal distribution (mean=155ms, standard deviation=25ms) and temporal extent ct drawn from a normal distribution (mean=21.66ms, standard deviation=4.17ms; the waveform was zero-padded outside a window duration of 6σ); the maximal amplitude of the N1 was drawn from a normal distribution (mean=2, standard deviation=1, arbitrary units). The randomly drawn P1 and N1 waveforms for each trial were summed, and then scaled by a multiplicative factor ERP_A_ or ERP_B_ dependent on the trial outcome A or B. Finally, this ERP was added to the white noise signal initially generated for that trial, to produce the final simulated electrophysiological signal.

In different simulations, the parameters n, f, t, mod, 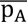, ERP_A_ and ERP_B_ were varied (sometimes jointly) to assess the robustness of the various phase opposition measures.

**Figure 2.**
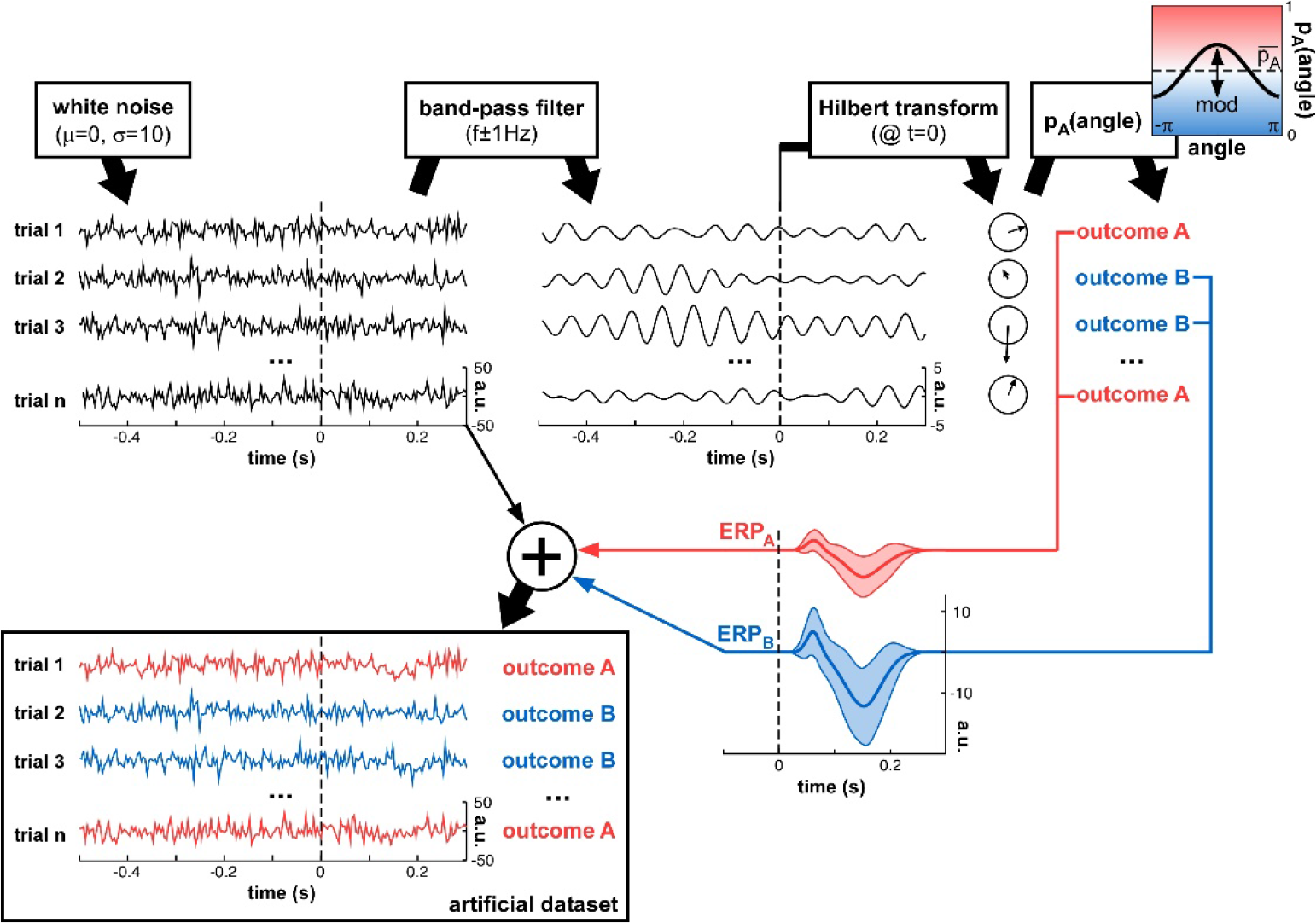
Artificial dataset creation. n trials were simulated, each with a (simulated) electrophysiological signal and an associated experimental outcome A or B. The signals were initialized with white noise, then band-pass filtered at a critical frequency f. The Hilbert transform extracted the oscillatory phase value at a critical time t. This phase was used to determine the experimental outcome for that trial: outcome A was randomly assigned with a mean probability 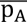 that was modulated as a cosine function of the phase angle. An ERP (with slightly randomized latency, duration and amplitudes for both P1 and N1 components, as illustrated by the shaded areas representing standard deviation across trials), whose overall amplitude depended on the trial outcome A or B, was added to the original white noise signal, to produce the final artificial dataset.

### 2.6 Test performance on artificial datasets: statistical power

The statistical power of a test is the proportion of experiments that would return a positive test result, given that an effect was indeed present. For a given set of parameters, we created 100 distinct artificial datasets, and then for each phase opposition measure, we evaluated the proportion of datasets yielding a successful statistical result. Based on the time t and frequency f of phase modulation implanted into the artificial datasets (see Figure 2), we defined a target time-frequency region (centered on point (t,f), with a tolerance of ±400ms and ±2 logarithmic frequency steps). An experiment was classified as a ‘hit’ if a statistically significant prestimulus phase opposition was detected *inside* the target region; a ‘false alarm’ occurred if significant prestimulus phase opposition was found *outside* of this region. The hit rate was thus defined as the proportion of datasets with a hit (out of 100 datasets), and the false alarm rate as the proportion of datasets with a false alarm. As previously (section 2.4.2, AUC), the hit and false alarm rates were contingent on the choice of statistical threshold. Thus, we varied the threshold systematically, and constructed the entire ROC curve; the statistical power was defined as the area under this ROC curve. Statistical power varies between 0 and 1. A statistical power value of 0.8, for example, would indicate that 80% of experiments could detect a significant phase opposition around the correct time and frequency (without also detecting it at incorrect times and/or frequencies); this value (or a higher one) is typically considered acceptable. Note also that chance level (the statistical power of a test with zero sensitivity) is not 0.5, but rather closer to 0.2 for most of our simulations. Indeed, since an ineffective test is equally likely to produce a significant result at any time-frequency point (inside or outside the target time-frequency region), the chance level can be simply evaluated as the ratio between the numbers of pre-stimulus time-frequency points lying inside vs. outside the target time-frequency region.

## 3. Results

### 3.1 Real experimental dataset (Busch et al, 2009)

First, it appears important to verify that all the statistical measures compared here share, at least, the ability to identify phase opposition in a real experimental situation, where the perceptual outcome is known to be modulated by ongoing oscillatory phase. To this end, we re-analyzed a previously published dataset (Busch et al., 2009). In that experiment, a brief peripheral target at detection threshold was presented on every trial, after a randomized inter-trial interval, such that observers reported only about half of the targets (hits) but missed the other half (misses). The phase bifurcation index (PBI) applied to this dataset (electrode Fz) had revealed significant phase opposition between these two trial groups at 7Hz, peaking 120ms before stimulus onset. We then explored the performance of other phase opposition measures on this same dataset (Figure 3).

**Figure 3.**
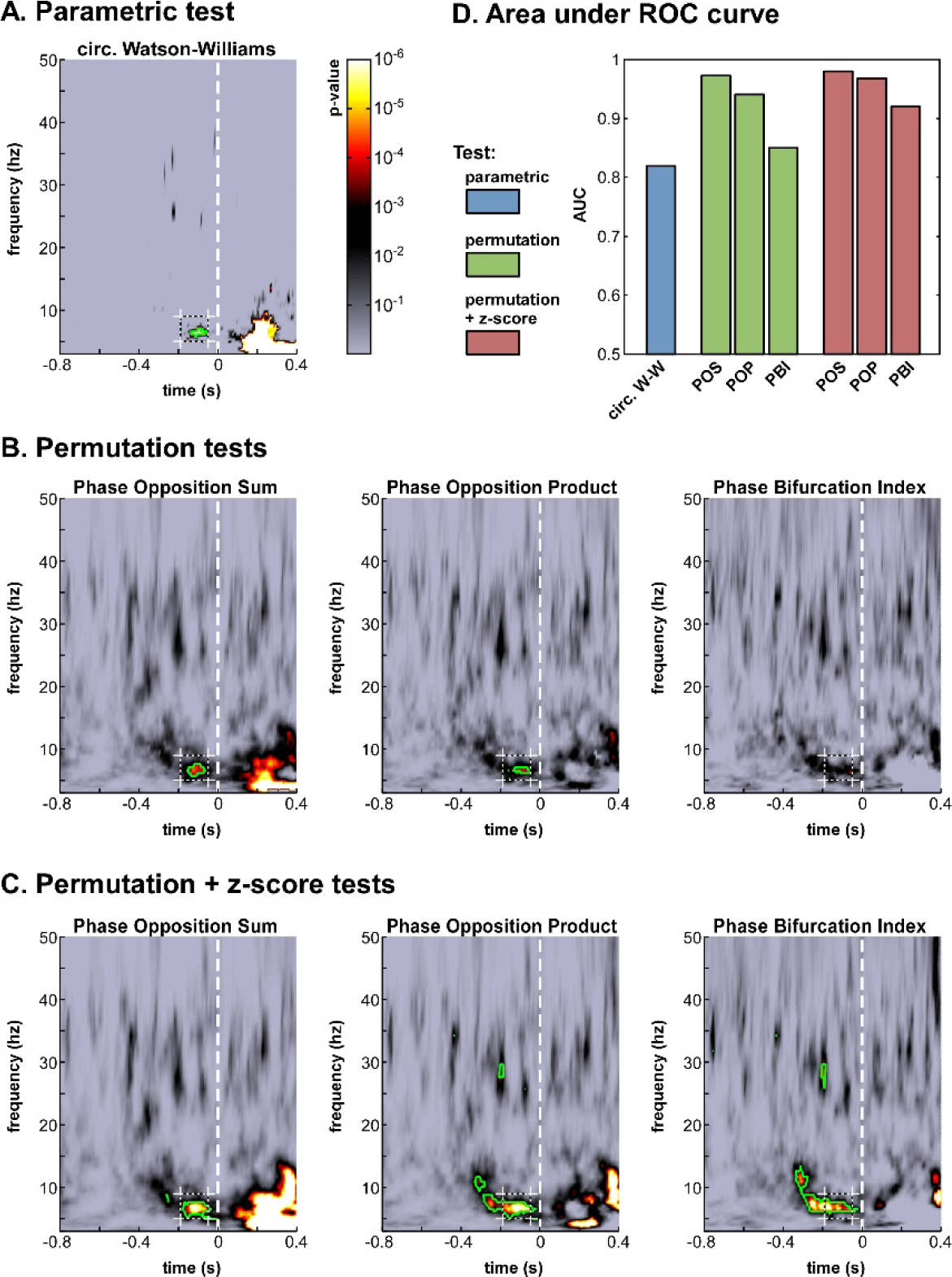
Phase opposition in a real-life experimental dataset. (Busch et al, 2009). **A-C**. Time-frequency maps of p-values on electrode Fz (colorbar in panel **A,** logarithmic scale) for the parametric test **(A),** the three permutation tests **(B)** and the hybrid tests combining permutation and z-score **(C).** The target time-frequency region (defined based on the results from the earlier study) is indicated by a dashed box. Time-frequency points with p-values below the threshold of FDR correction for multiple comparisons are outlined in green. **D.** Results of an AUC procedure contrasting the proportions of significant time-frequency points inside vs. outside the target time-frequency region. The bar colors correspond to those defined in **Figure 1**.

The seven time-frequency maps of p-values shown in Figure 3 (panels A-C) all demonstrated significant (p<0.0002) phase opposition around the same time (−120ms) and frequency (7Hz) as in the original study (Busch et al., 2009). For 6 of these 7 measures, the resulting p-values were robust to an FDR correction for multiple comparisons using alpha = 0.05 (Benjamini & Hochberg, 1995). It is worth noting that the last remaining measure was the PBI coupled with a standard permutation test (Figure 3B), precisely the one employed in the original paper. This indicates that our comparison did not unwittingly favor the initially applied measure. (In the original study, the PBI values were first combined across observers before applying the permutation test, resulting in slightly different p-values, which were in fact robust to FDR-correction; here, p-values were first obtained for each observer and subsequently combined across observers, for consistency with the procedure applied to the circular Watson-Williams test).

The different tests should not simply be compared in terms of their best p-values around the initially expected time-frequency point; indeed, a test’s efficiency also depends on its propensity to produce false positives, i.e., apparently significant phase opposition at a priori unexpected time-frequency points. To provide a comprehensive assessment of each test’s efficiency, we thus performed an ROC analysis, as follows. A target time-frequency region was defined based on the findings of the original study, centered on 7Hz and 120ms pre-stimulus, with a tolerance of ±2Hz and ±70ms. For a given statistical threshold, the ‘hit rate’ was defined as the proportion of time-frequency points *inside* the target time-frequency region with significant p-values (i.e., p-values below the statistical threshold), and the ‘false alarm rate’ as the proportion of pre-stimulus time-frequency points *outside* this region with significant p-values. After measuring hit rates and false alarm rates for all possible statistical thresholds, an ROC curve was obtained, expressing hit rate as a function of false alarm rate. The area under this ROC curve (AUC) can be used as a threshold-independent measure of a tests’ sensitivity.

We computed this AUC for all seven tests (Figure 3D). As could be expected from the p-value maps, all tests performed reliably, with AUC values above 0.8. The most sensitive test appeared to be POS, followed by POP and then PBI. This ranking was the same, whether a standard permutation test or a hybrid permutation+z-score test was applied, though the latter performed consistently (albeit marginally) better than the former. Finally, the circular Watson-Williams test yielded the lowest AUC value (0.82).^4^ Nonetheless, as will be seen later (section 3.2), the circular Watson-Williams test is often among the most sensitive measures of phase opposition.

### 3.2 Artificial datasets

The confirmation of the adequacy of all seven tests on a given ‘reference’ dataset was an essential step, but leaves open a number of questions. Would all tests prove equally robust to variations in specific properties of the dataset, such as the frequency of the rhythmic modulation, its magnitude, the number of trials collected, the presence and amplitude of post-stimulus ERPs, and so on? Answering these questions with real experimental datasets could easily require several hundred experiments, and a lifetime of data collection. To address these issues, we created instead a large number of artificial datasets, for which various parameters could be precisely controlled (Figure 2).

Each dataset included n trials with simulated electrophysiological signals (initialized as white noise) and an associated trial outcome A or B. The trial outcome was decided with a probability that was a sinusoidal function of the signal phase angle at a specific time and frequency. Finally, noisy ERP waveforms, of possibly different amplitudes for the two trial types A or B, were added to the original electrophysiological signal. The number of trials (n), the average likelihood of outcome A (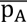), the depth (mod), frequency (f) and time (t) of phase modulation, as well as the average ERP amplitudes (ERP_A_ and ERP_B_) were parameters that could be varied in distinct simulations.

For each fixed set of parameters, we generated 100 random artificial datasets and applied the seven phase opposition measures. The performance of each measure was quantified by their statistical power: the proportion of datasets (out of 100) yielding a successful statistical outcome, with significant phase opposition detected inside but not outside a target time-frequency region. This statistical power was computed using a threshold-independent AUC procedure^5^. Statistical power values above 0.8 are conventionally considered appropriate. Therefore, a single value describing statistical power over the 100 simulated datasets can serve as a main criterion to assess the reliability of each measure (value>0.8). Nonetheless, for the additional purpose of comparing measures against each other, it is useful to keep in mind that differences of statistical power equal to or larger than 0.06 (e.g. 0.87 vs. 0.93) would normally be statistically significant at the p<0.05 level using a χ^2^ test.

#### 3.2.1 Depth of oscillatory phase modulation

The first set of simulations investigated the effect of the depth of oscillatory phase modulation (Figure 4). Indeed, this manipulation is directly linked to the detectability of phase opposition: with a phase modulation of 1 the outcome likelihood changes from 100% A to 100% B between one oscillatory phase and its opposite, and thus phase opposition should reach its maximal value; whereas with 0 phase modulation the likelihood of outcome A would be constant, and there would simply be no phase opposition to detect. For this first simulation, we arbitrarily fixed the other parameters as follows: number of trials n=500, frequency of phasic modulation f=15Hz, time of phasic modulation t=0 (i.e., the 15Hz phase value measured *at stimulus onset* influenced the trial outcome), equal likelihood of outcomes A and B 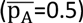, and no ERP produced after stimulus onset (while this last assumption is unrealistic, it will allow us to independently explore the influence of ERPs in later simulations).

**Figure 4.**
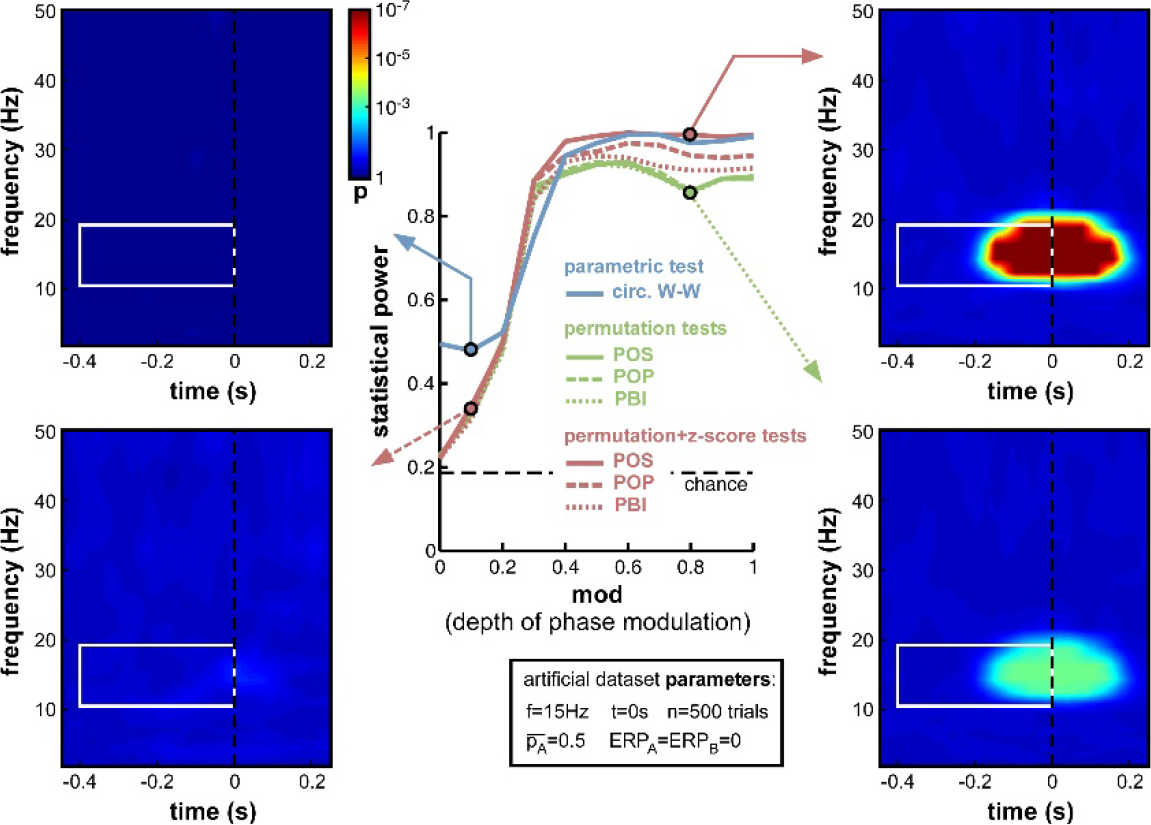
Effect of varying depth of oscillatory phase modulation. *The central plot illustrates the changes in statistical power caused by the parameter variations. The line colors correspond to those used in Figures 1 and 3. The solid blue line plots the statistical power of the circ. W-W test. Solid, dashed and dotted lines correspond respectively to POS, POP and PBI, in green for the standard permutation tests, and red for the hybrid permutation+z-score tests. The fixed parameters for these simulations are listed in the box below the main plot. In this and the following figures, four representative time-frequency maps of p-values (obtained by log-averaging the p-value maps from the 100 simulated datasets) are displayed for illustrative purposes. The arrow pointing to each map identifies the measure and the parameter value being illustrated. All maps share the same color scale (see colorbar next to the top-left map). The target time-frequency region is highlighted using a white box. Significant pre-stimulus time-frequency points inside vs. outside this box count as hits and false alarms (respectively) for the calculation of statistical power*.

As expected, upon decreasing the depth of modulation from 1 to 0, the statistical power of all seven measures went down (Figure 4), from high values (0.8 and above) to chance level (approximately 0.2, defined by the ratio of time-frequency points inside vs. outside the target region^6^). For all measures, the tipping point was around modulation depths of 0.3 to 0.4: above this point, all measures performed consistently well; below this point, the decrease in statistical power accelerated. This prompted us, in all subsequent simulations, to use a value of 0.4 for the ‘depth of modulation’ (mod) parameter, i.e., just above the “tipping point”: this still produced a satisfactory level of performance, yet also made this performance very susceptible to any deterioration induced by manipulations of the other parameters.

All seven measures behaved comparably in this first set of simulations, with minor differences in performance. At phase modulation values of 0.4 and above, both circ. W-W and POS (using the hybrid permutation+z-score test) virtually reached maximal statistical power (1), while POP and PBI (with the same hybrid test) reached respectively 0.97 and 0.94. The last three measures involving standard permutation tests (POS, POP, PBI) were systematically below the others, but still reached 0.92 performance. One reason why these tests might perform sub-optimally here could be an insufficient number of permutations, leading to insufficient precision of p-values. Indeed, only 1000 permutations had been used in these simulations. As can be seen in the time-frequency map on the bottom right of Figure 4, this implies that p-values from the permutation tests could not improve beyond 0.0005, even when the phasic modulation was maximal. Other tests, however, were not limited in this way, and could attain much more significant p-values (p<10^−7^, see top-right time-frequency map in Figure 4). Before continuing with other parameter manipulations, therefore, we decided to assess the importance of the number of permutations used in the comparison between tests.

#### 3.2.2 Number of permutations

While other parameters remained fixed as before, and the modulation depth parameter was set at mod=0.4, we varied the number of permutations from 10 to 10,000. Logically, this manipulation should only alter the non-parametric tests that rely on such permutations to estimate their null distribution. As expected, therefore, the parametric circ. W-W test was wholly unaffected by the variations (Figure 5). Although both types of non-parametric tests (permutation, or permutation+z-score) were expected to depend on permutation number, the hybrid permutation+z-score test appeared remarkably robust at low permutation numbers, with statistical power remaining above 0.9 even down to only 10 permutations (Figure 5). This suggests that the mean and standard deviation computed across only 10 surrogate (permuted) datasets are sufficiently precise estimates of the shape of the null distribution. In contrast, for the conventional permutation tests, statistical power was poor (around 0.6) with only 10 or 100 permutations, and kept improving from 1000 to 10,000 permutations (Figure 5). With that number, the permutation tests finally attained comparable statistical power to the hybrid tests.

**Figure 5.**
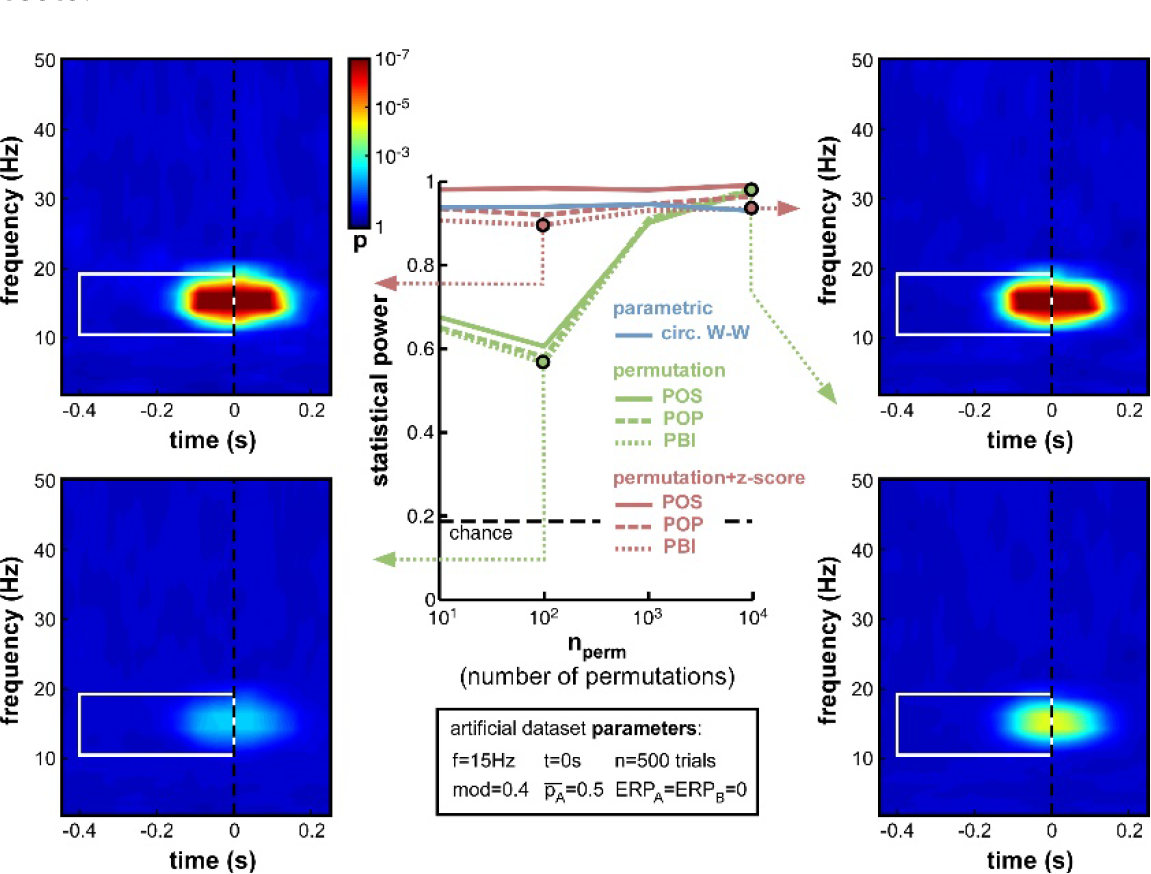
Effect of varying permutation number. Artificial dataset creation parameters are identical to those used in Figure 4, with the depth of oscillatory modulation set at mod=0.4. All four time-frequency maps illustrate the results of the PBI measure. Using a hybrid permutation+z-score test, PBI produces comparable results with 100 or 10,000 permutations (top-left and top-right). Using a standard permutation test, however, with 100 permutations the target p-values are hardly distinguishable from the background noise (bottom-left), whereas statistical power can be restored with 10,000 permutations (bottom-right).

The conclusions that can be drawn from this set of simulations are, therefore, that conventional permutation tests can be equivalent to the hybrid permutation+z-score tests, but large numbers of permutations (on the order of 10,000) are required to achieve this equivalence. Conversely, the hybrid tests can provide reliable estimates of the statistical power of a phase opposition measure, even when lower numbers of permutations are used. Consequently, both to simplify computations and to improve readability, in the following simulations we used only 1000 permutations, and shall not display anymore the results of the 3 conventional permutation tests^7^.

#### 3.2.3 Frequency of oscillatory phase modulation

A natural concern when measuring phase opposition is that the likelihood of detecting phasic modulations may depend on the frequency of the underlying oscillation. Intuitively, slower oscillations may be less prone to errors in phase measurement, simply because any fixed temporal uncertainty (say 10ms) in electrophysiological signals would translate into smaller phase variance at slower frequencies (only 1/10^th^ of a cycle at 10Hz, but 1/2 of a cycle at 50Hz). The results in Figure 6, however, demonstrate that oscillatory frequency has little influence on the statistical power of phase opposition measures under the conditions of these simulations. All measures remained over 0.8 as frequency varied from 7Hz to 40Hz. Across frequencies, the highest statistical power was given by POS, followed by POP, PBI and circ. W-W. Higher frequencies did not reduce statistical power, except for a marginal decrease at 40Hz concerning the circ. W-W test. This robustness is likely to be specific to the conditions of these simulations: ongoing oscillations were given equal power at all frequencies (i.e., created from white noise; see Footnote 2), and no ERP was included here that could have confounded phase measurements in specific frequency bands. (A later set of simulations, in section 3.2.7, will explore the effect of oscillatory frequency in the presence of sizeable ERPs.) Finally, the time-frequency maps in Figure 6 also underline that the temporal resolution of phase opposition measures, evident in the temporal spread of significant activations, is directly related to oscillatory frequency, with increased temporal precision at higher frequencies (reflecting, in part, the temporal extent of the wavelets used in the time-frequency transform).

**Figure 6.**
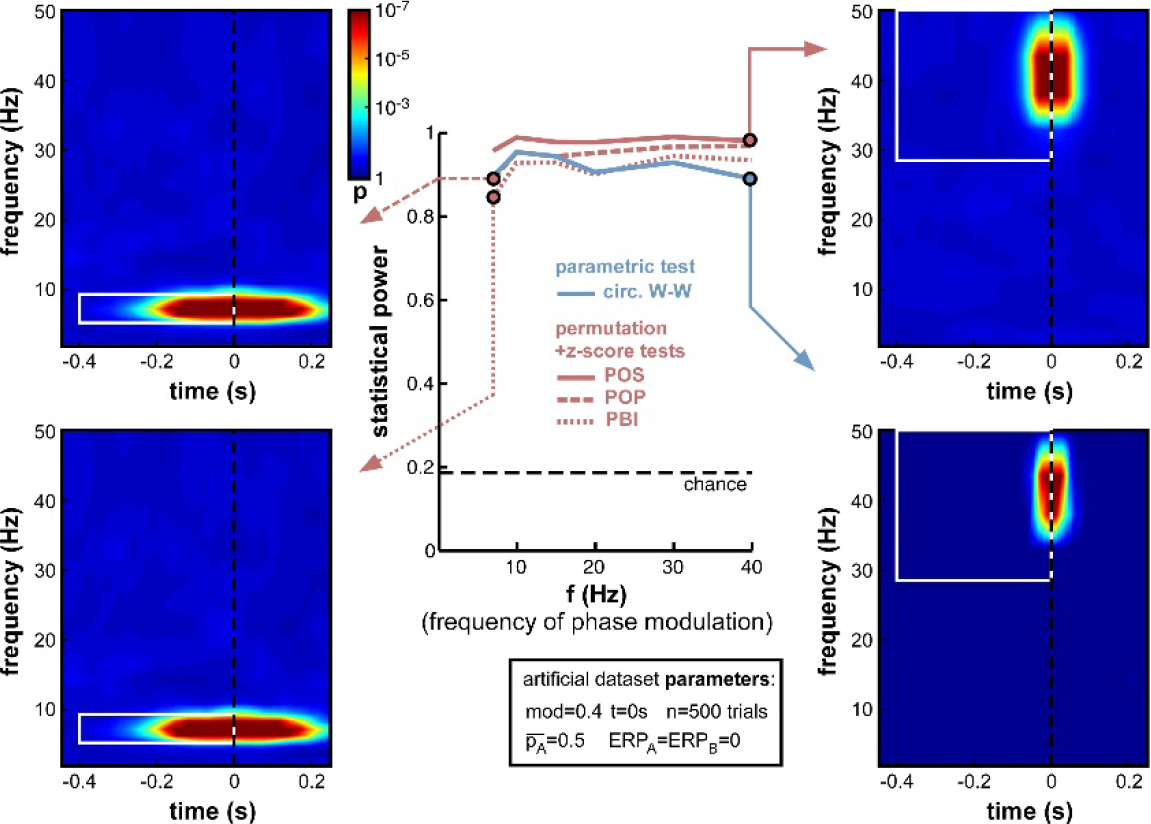
Effect of varying frequency of oscillatory phase modulation (in the absence of an ERP). For improved readability, only the parametric circ. W-W test and the hybrid permutation+z-score tests are presented in this and all subsequent figures. Each conventional permutation test (not shown) consistently mirrors the patterns of the corresponding hybrid test, only with marginally lower statistical power. All tests appear relatively robust to changes in oscillatory frequency.

#### 3.2.4 ERP amplitude

The onset of a sensory stimulus at time zero is normally followed by an event-related potential (ERP in EEG, also called event-related field or ERF in MEG), reflecting the activation of the sensory and perceptual apparatus of the corresponding modality, and potentially also a number of subsequent attentional and cognitive operations (Luck, 2014). ERPs have relatively high amplitude compared to that of ongoing brain oscillations. And, most importantly for our purposes, ERPs strongly affect measurements of oscillatory phase. It is still fiercely debated whether ERPs denote a phase reset of ongoing oscillations, or are merely superimposed on them (Makeig et al., 2002; Fell et al., 2004; Mazaheri & Jensen, 2006; Min et al., 2007; Sauseng et al., 2007); but in either case, phase measurements in time-frequency windows that overlap with ERPs cannot provide an independent estimate of ongoing oscillatory phase, and in some cases (depending on the existence of phase reset, and on the relative amplitude of ongoing and evoked signals) could be entirely dominated by the phase of ERP signals.

It appears necessary, therefore, to evaluate the statistical power of phase opposition measures in the presence of ERPs. Importantly, for these simulations, we assumed that the phase of background ongoing oscillatory signals, free of any ERP, was the key element in determining perceptual outcome (we had direct access to these background oscillatory signals since they had been artificially created; see Figure 2, top right). Subsequently, these background signals were summed with ERP waveforms, and the phase opposition measures were only given to operate on the summed signals (just as would happen in a genuine experimental situation). We could thus evaluate any disruption caused by ERPs masking the relevant phase of the background signal. In this set of simulations, we systematically varied the average ERP amplitude, but maintained it equal for the two perceptual outcomes, that is, ERP_A_=ERP_B_ (a later set of simulations in Section 3.2.9 will explore situations where the two ERPs are unequal).

**Figure 7.**
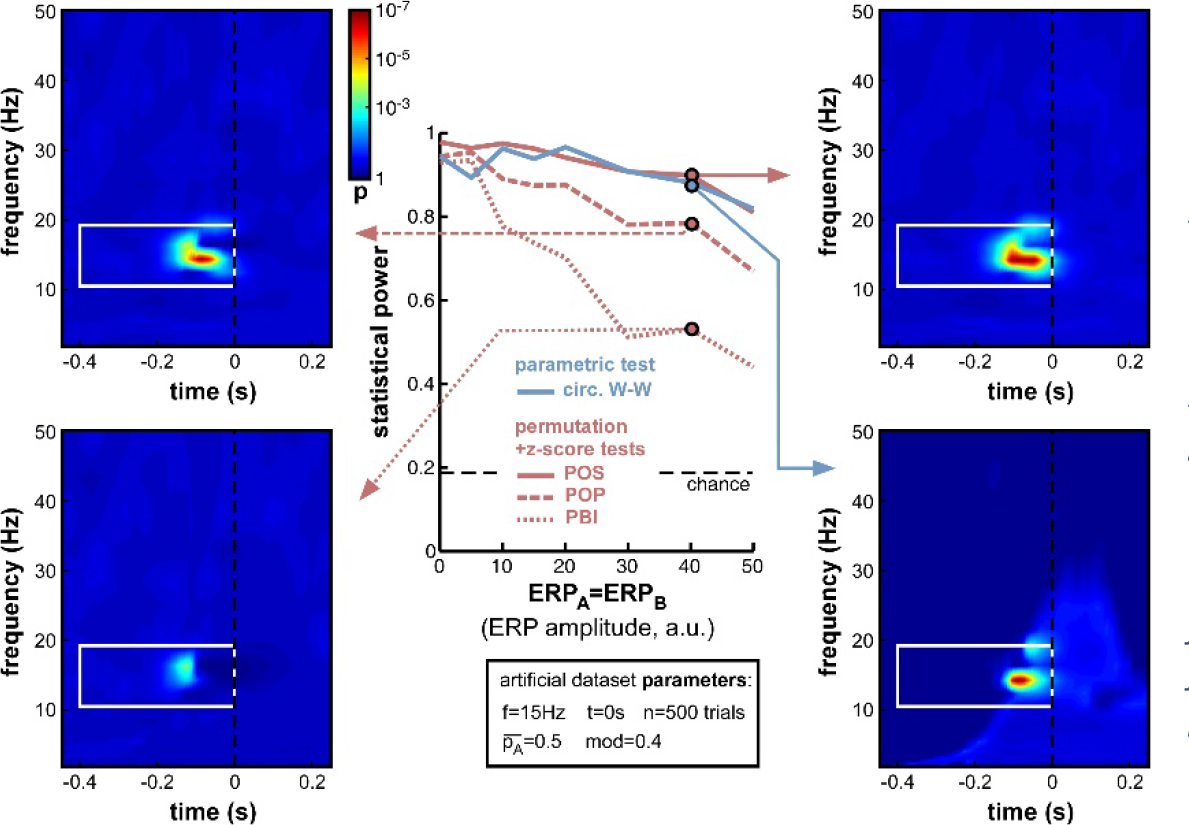
Effect of varying ERP amplitude. Parameters ERP_A_ and ERP_B_ were varied together from 0 to 50 (arbitrary units, the relevant point of comparison being the standard deviation of the background noise whose oscillatory phase was used to determine perceptual outcome: this standard deviation was fixed at 10 units in all simulations). As expected, the statistical power of all measures decreased with increasing ERP amplitude. Both POS and circ. W-W retained satisfactory power (above 0.8) at all amplitudes tested, while POP and PBI fell down to 0.67 and 0.44, respectively. Time-frequency maps illustrate distortions in the phase opposition landscape caused by ERPs, in particular a leftward shift of latency (compare, for example, to upper time-frequency maps in Figure 5). The time-frequency spectral signature of ERPs is visible as a lighter blue ‘hill’ in the bottom-right map.

As expected, increasing ERP amplitude had a detrimental effect on all phase opposition measures (Figure 7). POS and circ. W-W were the most robust measures, retaining statistical power above 0.8 for all ERP amplitudes tested, followed by POP and PBI, which decreased to 0.67 and 0.44 statistical power, respectively. The detrimental effect of ERPs on phase opposition is particularly visible in the four time-frequency maps of Figure 7 (compared, for example, to the upper two time-frequency maps of Figure 5). The region of significant phase opposition (when present) is smaller, and its center shifted to the left, i.e. 50-100ms earlier than the actual time t=0 at which phase modulation was applied. This shift can be understood as ERPs (whose phase is sensibly similar across all trials) “masking” the phase opposition between trial groups. The spectral signature of ERPs, and hence the time-frequency region where ERPs can potentially affect phase opposition, is clearly visible as a lighter blue ‘hill’ in the bottom-right time-frequency map.

A logical (albeit indirect) consequence of these simulations is that phase opposition may be less easily detected in experiments where stimulus onset is temporally predictable, because of e.g. a fixed inter-trial interval, or a preparatory cue (see also Footnote 1). Indeed, any pre-stimulus event bearing a fixed temporal relation to stimulus onset (such as the end of the preceding trial, or the onset of the preparatory cue) will evoke its own response activity pattern, affecting the phase distribution of “ongoing oscillations” in all trials, independently of the eventual trial outcome. This is more or less equivalent to the present simulations, in which a common “evoked” signal (an ERP) is affecting the phase of both trial groups in a similar manner, i.e. ERP_A_=ERP_B_ (with a major difference, of course, in the time at which this influence is exerted–early vs. late in the pre-stimulus interval).

For all subsequent simulations (except the further investigations of ERPs in Section 3.2.9), we shall fix the ERP amplitude parameters to ERP_A_=ERP_B_=10, an intermediate value equivalent to the amplitude of the background noise (whose oscillatory phase serves to determine the trial outcome, see Figure 2, top-right).

#### 3.2.5 Trial number

The reliability of inter-trial coherence estimates is notoriously dependent on trial number (Moratti, Clementz, Gao, Ortiz, & Keil, 2007; Vinck, van Wingerden, Womelsdorf, Fries, & Pennartz, 2010). Phase opposition measures directly inherit this dependence (Eqs (4–7)). As trial number is a key element of any experimental design, which can limit or even forbid addressing certain experimental questions, the exact influence of this parameter (n) is important to assess.

When decreasing trial number n from 1000 down to 10 (Figure 8), the anticipated decrease of statistical power occurred for all measures, yet at slightly different rates. Non-parametric measures decreased slowly down to 200 trials, and faster afterwards (reaching levels close to chance at 50 trials and below). POS, for example, which showed the highest statistical power (0.985) at 1000 trials, remained above 0.8 down to nearly 200 trials. In contrast, the parametric measure (circ. W-W) decreased much faster. Even though it was on par with POS down to 500 trials, at 200 trials and below it had much lower statistical power than all other measures. At 100 trials it showed no significant phase opposition (chance level=0.19; bottom-left time-frequency map), while POS could still detect phase opposition in more than half of the simulations (0.56; top-left time-frequency map).

Another lesson that can be learned from this set of simulations is that, whatever the measure used (even PBI, which so far systematically displayed the worst statistical power), a reliable estimate of phase opposition can always be obtained given enough trials (on the order of 1000 trials in these simulations; Figure 8).

**Figure 8.**
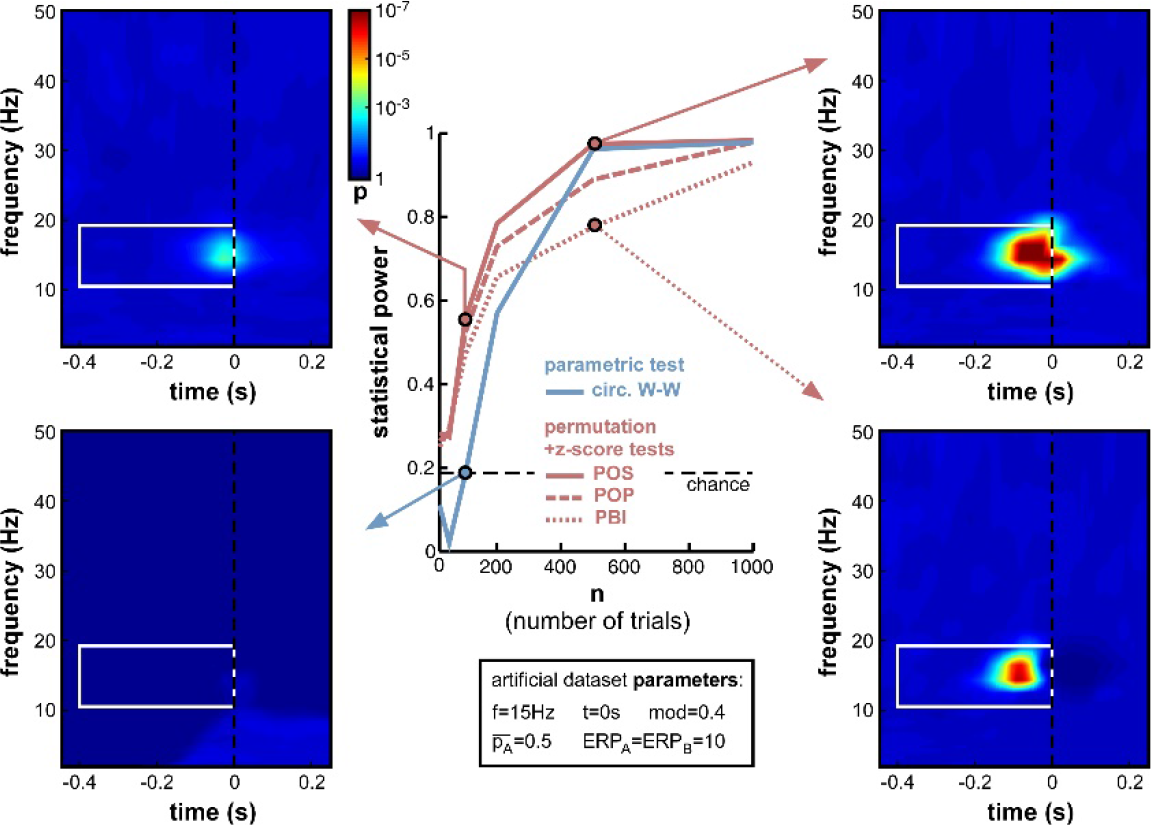
Effect of varying trial number. Parameter values used for n were 10, 50,100, 200, 500 and 1000. While all measures were comparably efficient using 1000 trials, their statistical power systematically decreased when less trials were simulated. Below 500 trials, circ. W-W decreased much faster than other measures, reaching chance performance at 100 trials (bottom-left time-frequency map), while other measures still showed above-chance performance (top-left time-frequency map).

#### 3.2.6 Relative trial number

Not only the total number of trials n, but also the relative number of trials in each group can influence phase opposition measures. In the previous simulations the experimental outcomes A and B were always equiprobable 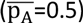. But experimental designs can often be biased (voluntarily or involuntarily) such that one outcome is more likely than the other. For example, the proportion of correct trials in a two-alternative forced-choice discrimination task should always be well above 50% (chance level); any phase opposition between correct and incorrect perceptual outcomes in such a task would thus be plagued by different trial numbers for the two groups. In turn, this difference may influence measures of phase opposition, either because one of the two groups would count an insufficient number of trials (as explored in the previous simulations, section 3.2.5), and/or because the reference quantity ITC_all_ would be biased towards ITC_A_ or ITC_B_ (which would consequently affect all phase opposition measures as described in Equations (4–7)).

Indeed, this new set of simulations revealed that all phase opposition measures were only truly efficient when outcomes were equiprobable, i.e. 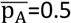 (Figure 9). Any departure from this equilibrium was sanctioned by a loss of statistical power, most measures falling below 0.8 if one of the outcomes was twice as likely as the other (i.e., 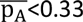 or 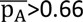). The most robust measure against this parameter manipulation was circ. WW, followed by POS, then by POP and PBI.

Interestingly, a strong asymmetry was apparent in the statistical power curves of the circ. W-W and POS measures (Figure 9), with lower performance when A was more likely than B (i.e. in the second half of the curves). Circ. W-W retained 0.78 statistical power with 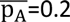 (see top-left time-frequency map), but fell down to 0.43 with 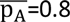 (see bottom-right time-frequency map). This puzzling result (considering that A and B were arbitrarily assigned labels) can in fact be explained by (Equation (9): due to the cosine function, outcome A was always maximal around phase 0, and outcome B at the opposite phase. This arbitrary choice effectively broke the symmetry of the dataset when in the presence of ERPs: the phase of the ERP signal around the critical time, similar for the two trial groups, could help (top-left map) or impair phase opposition detection (bottom-right map),depending on which of the two trial groups had a phase closest to that of the ERP^8^.

Overall, the strong decrease of statistical power obtained with unbalanced datasets (Figure 9) has important implications for experimental design and data analyses practices. Equiprobable outcomes should be sought whenever possible. Of course, a dataset resulting from an unbalanced experimental design can always be subsampled to restore equal numbers of trials in the two groups (e.g. by randomly rejecting trials from the group having higher likelihood), but this will necessarily come at the cost of decreasing the overall trial number, which, as demonstrated above (Figure 8), can also have potentially drastic consequences.

**Figure 9.**
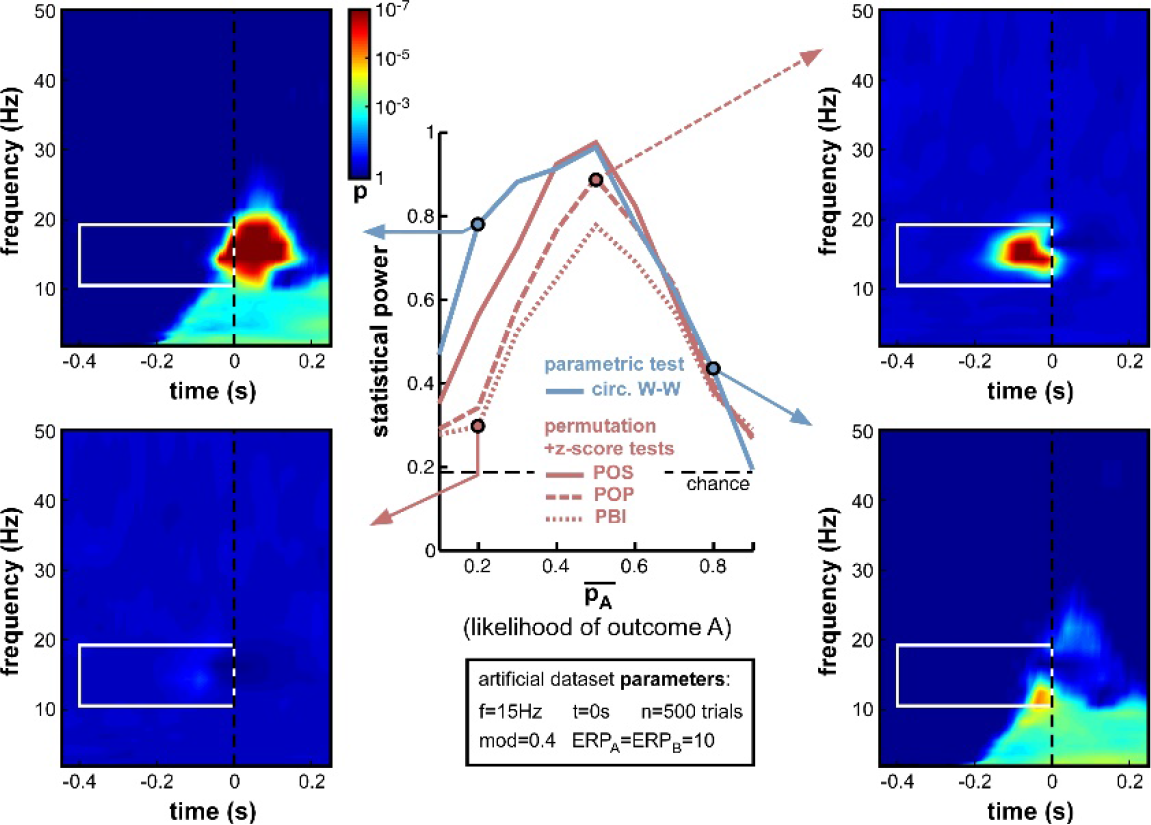
Effect of varying relative trial number. (likelihood of outcome A). Parameter 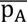 was varied from 0.1 to 0.9 in steps of 0.1. Optimal performance was obtained in a situation with equiprobable outcomes (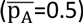), while unbalanced situations strongly impaired statistical power. Circ. W-W and POS were the most successful tests, followed by POP and PBI. The asymmetry that is particularly visible for the circ. W-W test (compare top-left to bottom-right time-frequency maps) is caused by an interaction between the phase of ERPs and the ongoing oscillatory phase that favors outcome A vs. B.

#### 3.2.7 Frequency of oscillatory phase modulation (with ERP)

The presence of an ERP, contaminating time-frequency estimates around stimulus onset and from roughly 0 to 30Hz (in our simulations), impedes the statistical power of phase opposition measures when the phase modulation is applied at t=0 and for 15Hz ongoing oscillations (Figure 7). It is likely, however, that this contamination might differ were phase opposition applied for a different frequency or at a different time. This set of simulations and the next were designed to address these two questions.

First, we varied the oscillatory frequency f at which the phase modulation was applied, exactly as in Figure 6, but this time in the presence of a sizeable ERP (ERP_A_=ERP_B_=10). The results revealed the same pattern as in Figure 6, only exacerbated (Figure 10). POS was the most successful measure at nearly all frequencies. Circ. WW performed well at low frequencies (below 15Hz, see top-left time-frequency map), but was the least successful above 20Hz (down to 0.7 statistical power at 40Hz, see bottom-right time-frequency map). POP and especially PBI displayed the opposite behavior, strongly affected by ERPs at low frequencies (PBI reaching down to 0.69 statistical power at 7Hz, see bottom-left time-frequency map^9^, yet equivalent to the best measure (POP) at frequencies above 30Hz (see top-right time-frequency map).

**Figure 10.**
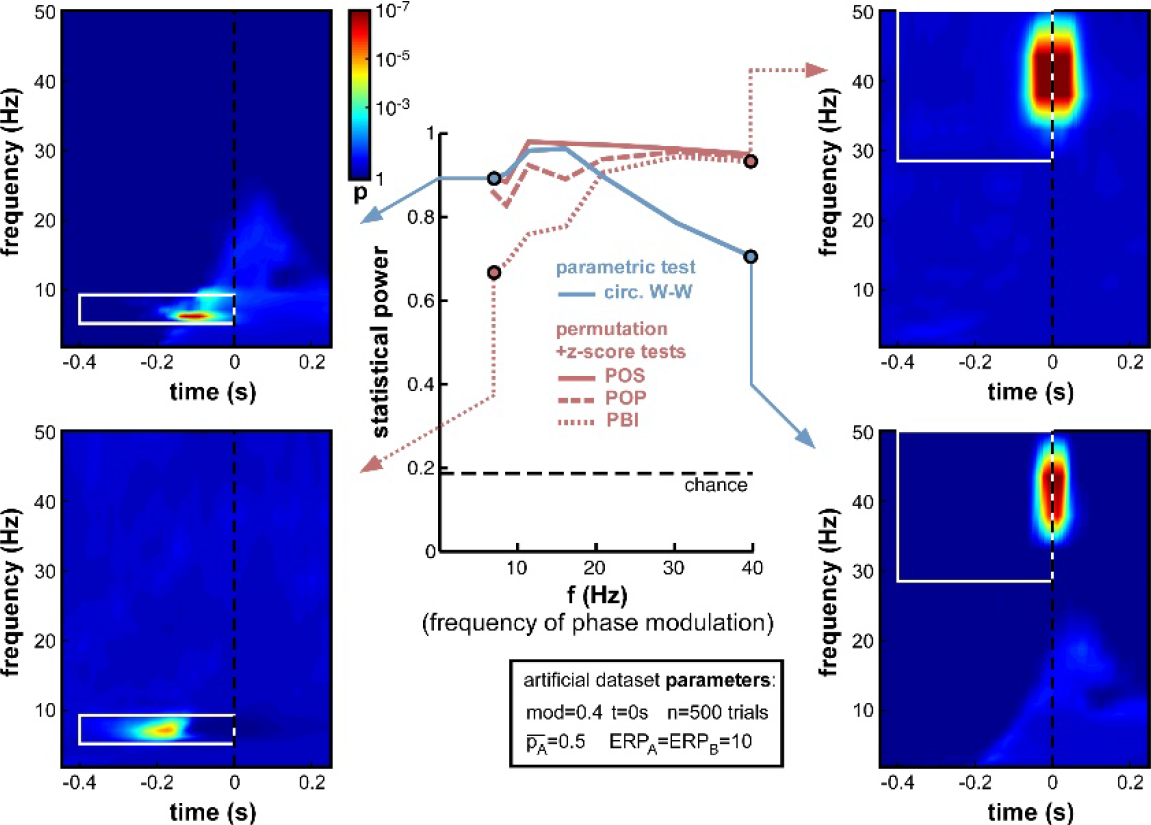
Effect of varying frequency of oscillatory phase modulation in the presence of an ERP. The pattern of frequency-dependence for the different phase opposition measures is similar to that observed in Figure 6, only exacerbated by the presence of an ERP. POS was consistently the highest-performing measure. Circ. W-W performed best at low frequencies (top-left time-frequency map) and worst at high frequencies (bottom-right map). PBI (and, to a lesser extent, POP) showed the opposite pattern, with maximal performance at high frequencies (top-right map) and minimal performance towards low frequencies (bottom-left map).

#### 3.2.8 Time of oscillatory phase modulation

Second, we varied the time t at which phase modulation was applied, from −200ms to +100ms relative to stimulus onset (the frequency f was maintained at 15Hz for this set of simulations). If a sensory or cognitive function operates periodically under the effect of an ongoing brain oscillation, the moment at which oscillatory phase should maximally influence the outcome of this function (and thus, phase opposition should be maximal) is the precise moment at which this function comes into play. In all logic, this should happen after stimulus onset, around the activation latency of the corresponding brain region or network. Unfortunately, this moment should also coincide with the emergence of ERPs, which have the power to mask phase opposition (Figure 7).

Indeed, the simulations (Figure 11) confirmed that, while all measures performed at ceiling (statistical power > 0.9) to detect phase opposition that was introduced at early latencies (at or before-50ms pre-stimulus), their statistical power dropped to inadequate values (<0.5) by 100ms post-stimulus, i.e. right in the middle of the ERPs. PBI plunged first, followed by POP, circ. W-W and finally POS. These last two measures were still remarkably efficient (power > 0.95) at the exact time of stimulus onset (t=0).

It is worth insisting that our calculation of statistical power only took into account pre-stimulus time-frequency points, even in simulations where the phase opposition was actually applied at post-stimulus latencies. This choice is justified by two reasons. First, the stationarity of oscillatory signals and the window length of the filtering operations required to extract phase values should imply that information about the critical oscillatory phase is already available in the pre-stimulus time window (even when the critical time t is itself in the poststimulus period). Second, post-stimulus “phase opposition” may be spuriously introduced by the ERPs, as illustrated for example in the bottom-right time-frequency map of Figure 9 concerning circ. W-W, or in the bottom-left map of Figure 12 concerning POS. In these maps, large time-frequency regions of (spurious) “significant phase opposition” can be observed in the post-stimulus region and at low frequencies (below 10Hz), even though no phase modulation actually occurred in this time-frequency range. We took the point of view of an experimenter who does not have access to the ground truth, and must therefore beware that any post-stimulus effect of phase could be the product of ERP-induced artifacts; and we thus restricted our search for phase opposition to the pre-stimulus time window^10^.

In sum, this set of simulations indicated that phase opposition becomes increasingly difficult to detect if it occurs in a time window coinciding with strong ERPs, and contrarily, increasingly easy to detect when it occurs at early latencies. Of course, this latency is not under the experimenter’s control, but depends on the sensory or cognitive function under study. In practice, this implies that phase opposition experiments may be more likely to succeed when they investigate the periodicity of a “peripheral” brain function (with short activation latency) rather than a more “central” one (with later activation latency). In any case, POS and circ. W-W tests provide the best outcome in such situations.

**Figure 11.**
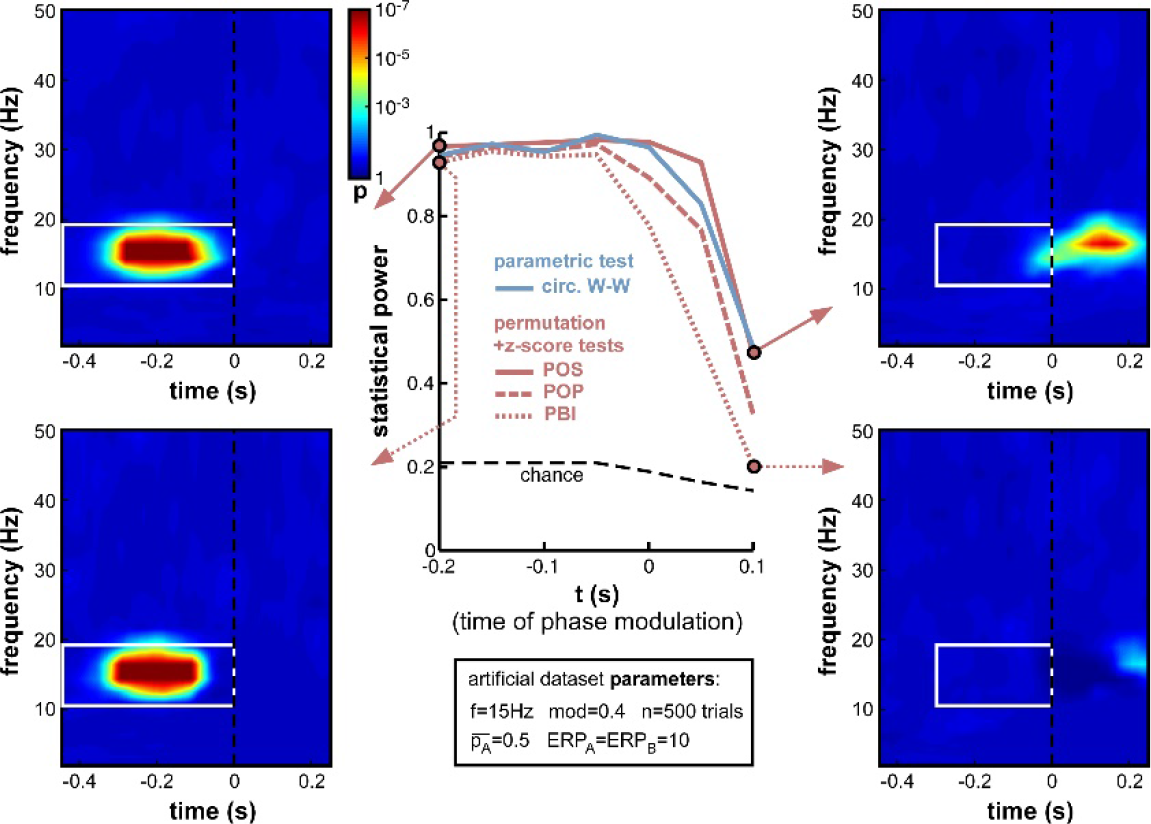
Effect of varying time of oscillatory phase modulation. The phase that determined the trial outcome A vs. B was extracted from background electrophysiological signals (by filtering and Hilbert transform, see Figure 2) at different moments around stimulus onset. The presence of an ERP with strong phase-locking across (all) trials may be expected to obscure the underlying phase opposition at specific time-frequency points. For latencies between −200 and −50ms, all phase opposition measures performed optimally (statistical power>0.9). With increasing latencies, all measures deteriorated, in the following order: PBI first, followed by POP, circ. W-W and finally, POS.

#### 3.2.9 ERP amplitude difference

So far, the influence of ERPs on phase opposition has only been investigated in situations where both task outcomes A and B produced ERPs of similar amplitude (ERP_A_=ERP_B_). This influence was found to be already fairly detrimental in such situations (see, in particular, Figures 7 and 11). But many experimental tasks can also be expected to produce different ERPs for different task outcomes. For example, the amplitude of certain ERP components is commonly found to vary, depending on the subject’s behavior. In one extreme situation, with brief visual stimuli at detection threshold, Busch et al (2009) found a strong ERP for hits but virtually no ERP amplitude for misses. As the potential masking of ongoing oscillatory signals by ERPs is directly related to ERP amplitude (whether this amplitude is the reflection of a phase reset process and/or of an additive signal), an asymmetry in ERP amplitudes for the two trial groups A and B could translate into an imbalance between the quantities ITC_A_ and ITC_B_ (relative to their baseline ITC_all_), and thus affect the calculation of phase opposition (through (Equation 4–7).

To explore the effects of such an ERP imbalance^11^, we fixed the ERP amplitude of trial group B to ERP_B_=10 (arbitrary units), and systematically varied the amplitude ERP_A_ of the other trial group from 0 to 50. The resulting statistical power curves (Figure 12) revealed that ERP imbalance could indeed impair phase opposition: all curves peaked around ERP_A_=10, that is, when the two ERPs were approximately equal. In fact, the performance decrease appeared nearly proportional to the ratio between highest and lowest ERP amplitude, regardless of whether ERP_A_ was above or below ERP_B_. This means that the impairment due to ERP amplitude imbalance cannot be compensated for by one of the two trial groups having low (or even null) ERP amplitude, and thus low contamination of phase calculations. PBI was overall the least successful measure, followed by POP. While POS performed above circ. W-W (and all other measures) around the amplitude equilibrium point (ERP_A_ between 5 and 20), this relation reversed and circ. W-W became the most successful measure when imbalance increased. In fact, POS performed worse than any other measure with ERP_A_≤2, and worse than circ. W-W and POP (but not PBI) with ERP_A_≥40.

**Figure 12.**
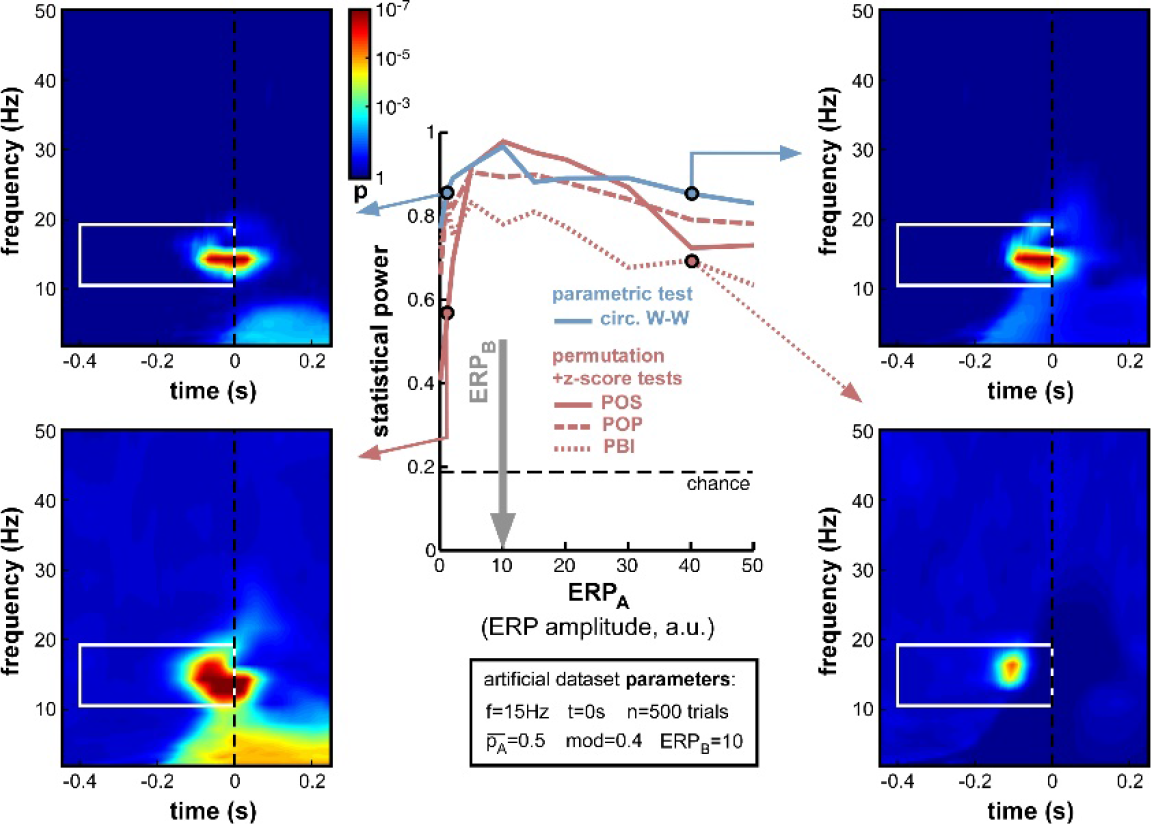
Effect of varying ERP amplitude difference. ERP amplitude for one group of trials (group B) was fixed at ERP_B_=10 arbitrary units (indicated by the vertical gray arrow). For all phase opposition measures, statistical power was best when ERP_A_ was approximately equal to ERP_B_, and declined both when ERP_A_ was higher (right time-frequency maps) and when it was lower (left maps; even though the phase of group A trials could be more accurately estimated in this latter case). When ERP_A_ and ERP_B_ were approximately equal, POS was the best performing measure. However, the POS measure became less reliable than others when the amplitude ratio was higher than 2 or lower than 0.5 (bottom-left map). In that case, the circ. W-W test provided the highest statistical power (top time-frequency maps).

To summarize this set of simulations, while it is best not to have any ERP contamination (see Figure 7), whenever ERPs are present (i.e., in most situations) it is better to have them as equal as possible for the two trial groups. Around the equilibrium point, POS has the highest statistical power, but when the ERP amplitude ratio is above 2 (or below 0.5), circ. W-W becomes preferable.

## 4 Discussion

Seven measures of phase opposition were compared in a variety of real and artificial experimental situations. All measures proved their value on a previously published dataset (Busch et al, 2009; Figure 3). On a series of artificial datasets with systematic manipulations of experimental parameters, we were able to provide a detailed assessment of all methods, and understand their common or individual reactions to changes in certain key parameters. These results informed us about the conditions in which phase opposition experiments are most likely to succeed, and about the right measure(s) to use for every situation.

### 4.1 Limitations

The present explorations, while hopefully useful, are necessarily incomplete. Other parameters or parameter combinations would certainly have been interesting to test. For example, the influence of ERPs (Figure 7) or of the timing of phasic modulation (Figure 11) are likely to be frequency-dependent, that is, different patterns of results might have been obtained if a slower (say, 7Hz) or a faster (say, 30Hz) phasic modulation frequency had been employed. In addition, changes in the shape and/or latency of ERPs between trial groups (which were not included here) could be expected to influence the statistical power of our different measures by introducing spurious phase opposition. Changes in the noise structure (e.g. from white noise to pink or brown noise; see Footnote 2) are also likely to affect the frequency-dependence of phase opposition (Figures 6 and 10). The range of possible factors to consider is virtually limitless. Nonetheless, we can hope that the key conclusions obtained in the various simulations performed here could generalize to a larger range of investigations.

Similarly, the choice of seven particular phase opposition measures necessarily leaves aside a number of other alternatives. Circular-to-linear correlation can be employed, for example, in experiments with a graded task outcome (e.g. reaction time), which would have been grouped here into two outcomes A and B (e.g. faster vs. slower reaction times). Studies that are motivated by a strong a priori about the relevant oscillatory rhythm (e.g. occipital alpha) could dispense with time-frequency phase opposition analyses altogether (comparing phase distributions across task outcomes), and instead plot task outcome as a function of phase (similar to the top-right plot in figure 2) in order to evaluate the depth of any sinusoidal modulation. Besides, standard intertrial coherence in (Equation 4–7) could be replaced by an equivalent but unbiased measure of phase-locking such as the pairwise phase consistency or ‘PPC’ (Vinck et al., 2010). Causal filters (Zoefel & Heil, 2013) or ERP removal and interpolation procedures (Lakatos et al., 2013; Henry et al., 2014) could be employed to avoid post-stimulus contamination of phase measurements. By no means do we wish to imply that the measures tested here are the only alternatives; however, all these measures have been employed recently by our group and others worldwide to explore phase opposition in time-frequency electrophysiological data. We thus anticipate that the comparison of these procedures should prove helpful to at least these experimenters, and hopefully also to others that may follow the same inspiration.

### 4.2 Key messages

For all three ad-hoc phase opposition measures introduced here (POS, POP, PBI), a streamlined non-parametric statistical procedure using a relatively small number of surrogates (permutations) to characterize the mean and standard deviation of the null hypothesis distribution, and subsequently deriving p-values from a z-score, was found to be preferable to the standard non-parametric permutation procedure. With large numbers of permutations, the two statistical procedures were equally efficient (Figure 5), but contrary to the standard procedure, the hybrid permutation+z-score tests did not suffer noticeably when the permutation number was decreased. With 1,000 permutations, as used in most of our simulations, this hybrid permutation+z-score test always outperformed the conventional permutation procedure. We thus recommend using the hybrid technique, or else using the conventional permutation method with sufficiently large numbers of permutations (at least 10,000).

No single measure was systematically better than all the other ones; rather, the optimal measure was strongly tied to the exact parameter values of the artificial dataset generation process. Across all simulations (Figures 4-12), however, the winning measure was *always* either POS or circ. W-W, and *never* POP or PBI. We thus recommend abandoning these two measures, especially PBI which we had introduced in 2009 (Busch et al., 2009) and which has been independently employed in (at least) 13 publications since (Hamm et al., 2012; Ng et al., 2012; Auksztulewicz & Blankenburg, 2013; Hanslmayr et al., 2013; Manasseh et al., 2013; Rana et al., 2013; Diederich et al., 2014; Park et al., 2014; Li et al., 2015; Shou & Ding, 2015; Strauss et al., 2015; van Diepen et al., 2015; Batterink et al., 2016). Although there is no indication that PBI would have led authors to erroneous conclusions in any of these studies, it is also clear that more accurate results (or higher statistical power) would have been achieved using other measures.

Under what conditions should the non-parametric POS measure be used, and under what conditions the parametric circ. W-W test? In many of the situations tested, both of these did provide equivalent results (and it might then be preferable to use the parametric method, if only because it does not require computationally intensive permutations). In specific cases, however, their performance was found to differ. POS was reliably better with low trial numbers (200 or less; Figure 8), for higher frequencies of phase modulation (20Hz and above; Figures 6 and 10), and for phasic modulations occurring just after stimulus onset (around 50ms; Figure 11). In turn, circ. W-W was better in asymmetric situations where either the relative trial number (Figure 9) or the ERP amplitude (Figure 12) differed markedly between the two trial groups. As many of the relevant parameter values cannot be known in advance and are not under the experimenter’s control (e.g. frequency or timing of phasic modulation), we recommend systematically using both methods. In order to make these analysis methods more easily accessible to all experimenters, and to make the results more easily comparable between studies, we provide <u>associated Matlab code</u> (http://www.cerco.ups-tlse.fr/~rufin/PhaseOppositionCode/) to automatically compute p-values derived from circ. W-W and POS procedures (with both standard permutation and hybrid permutation+z-score tests implemented for the latter).

### 4.3 Conclusion

Independent of the exact analysis procedure employed, our explorations have revealed that the likelihood for an experimenter to detect phase opposition in an electrophysiological dataset is affected by several factors. This was true, even using a constant magnitude of phasic modulation (for example, in most of our simulations a reversal of rhythmic phase produced a 40% modulation of the probability of task outcome A vs. B; Figures 5-12). Among the most important factors to consider for phase opposition analyses are the absolute and relative trial numbers (Figures 8 and 9), the overall ERP amplitude (Figure 7) and any potential ERP amplitude difference between the trial groups (Figure 12), as well as the frequency (Figure 10) and exact time (Figure 11) of the phasic modulation. Throughout all these simulations, the phase bifurcation (PBI) measure performed systematically worse than others. The optimal statistical power was shared (alternately) by one parametric measure, the circular Watson-Williams test (circ. W-W) and a non-parametric one, the phase opposition sum (POS). Our conclusion is, therefore, to recommend using these two tests in any future exploration of phase opposition. The analysis code that we provide should hopefully facilitate that objective.

## 5. Acknowledgments

This work was supported by an ERC Consolidator grant P-CYCLES number 614244. The author wishes to thank all past and present members of his lab for useful discussions on phase opposition, and in particular D. Lozano-Soldevilla, S. Crouzet and B. Han for further providing helpful comments on the manuscript.

## Footnotes

1 Although we do not explicitly explore alternative experimental situations with temporally predictable stimulus onset (caused e.g. by a fixed inter-trial interval, or by a preparatory cue), the behavior of phase opposition measures in such situations can be inferred somewhat from our simulations with outcome-independent ERPs (see Figures 7 and 11). Indeed, such a post-stimulus ERP would have a similar influence on ongoing oscillations as would a pre-stimulus locking signal (e.g., the end of the previous trial or preparatory cue), except that the latters influence would be visible at a much earlier time, in the pre-stimulus window.

2 White noise comprises equal power at all frequencies. In contrast, electrophysiological signals generally display higher power towards lower frequencies, e.g. pink or brown noise (depending on the logarithmic exponent of the frequency/power relation). The choice of white noise here was only a first approximation, to simplify comparisons of phase opposition measures across frequencies (since all frequencies have a priori equivalent signal-to-noise ratio) and to avoid arbitrarily deciding on a specific logarithmic exponent (whose exact value can depend on several experimental factors).

3 The choice of a cosine function, with maximal likelihood of outcome A at phase 0, was arbitrary. This choice would not be expected to affect the results of any simulation (except as described in Figure 9). Generally, as explained also in Section 2.1, the nature of EEG/MEG recordings is such that the polarity of oscillations detected at the scalp is only indirectly related to the underlying cortical oscillations. Consequently, absolute EEG/MEG phase cannot be easily interpreted, and we focus instead on relative phase differences between the two trial groups.

4 This apparently lower performance is likely to stem from the all-or-none nature of this parametric test, coupled with the stringent demands of our ROC analysis. 95% of the p-values from this test were higher than 0.999 or lower than 0.001. Within the target time-frequency region, 31% of time-frequency points had the highest possible p-value (compared against 88% outside the target region). Thus, as the statistical threshold grew progressively more liberal, the hit rate gradually increased to a value of 0.69 (i.e., 69% of time-frequency points inside the target region gave significant p-values), while the false alarm rate remained below 0.12; however, with the next increase of statistical threshold, both hit rate and false alarm rate suddenly reached 1, and the ROC curve was thereby truncated.

5 Different from the previous analyses (Figure 3), in which only one dataset was available, here the hit and false alarm rates were not defined as proportions of time-frequency points, but as proportions of datasets (out of 100). This implies that a single significant p-value inside the target time-frequency region of all 100 datasets could suffice to yield 100% hit rate, and thus, that the all-or-none nature of the circular Watson-Williams test was less likely to impair its AUC performance here.

6 Except for the circular Watson-Williams test, which levelled off around 0.5 statistical power because the likelihood of significant p-values inside vs. outside the target region was exactly the same for this test, namely zero.

7 The 3 permutation tests were still performed, however; the resulting data (not shown) consistently reflected the aforementioned conclusion: statistical power was systematically (albeit marginally) lower than in the corresponding hybrid tests, and followed the same overall pattern.

8 To verify this, another set of simulations (not shown here) was performed after changing the + sign into a −sign in Equation (9). This effectively changed the optimal phase for outcome A, from zero to pi radians. The results showed again an asymmetry, but this time with higher performance when A was more likely than B.

9 This map also best illustrates a point already made earlier (Figure 7), that the presence of an ERP can strongly distort the time-frequency landscape of phase opposition. This is particularly true at lower frequencies. Here, at 7Hz, the peak of phase opposition is registered about 200ms earlier than the true time of phase modulation (t=0). In fact, these simulations help explain why many previous studies of phase opposition have reported effects peaking well before stimulus onset (from −100ms to −300ms or even earlier), when logic dictates that the critical time of phase opposition should be around or just after stimulus onset (see Section 3.2.8).

10 In fact, restricting analyses to pre-stimulus time points still does not fully guarantee that observed effects are devoid of ERP contamination. Especially at low frequencies, when the half-length of the temporal window used for time-frequency decomposition exceeds the latency of ERPs, ERP contamination can spread to pre-stimulus time points. For example, the window of potential contamination for our simulated ERPs is visible as a lighter-blue region in the time-frequency map at the bottom-right of Figure 7. To rule out this possible confound, several studies have explicitly verified that significant pre-stimulus phase opposition effects were detected outside of this “contamination window” (e.g., Busch et al, 2009). Other strategies involve using causal filters (Zoefel & Heil, 2013) or ERP removal and interpolation techniques (Lakatos et al., 2013; Henry, Herrmann, & Obleser, 2014).

11 We focused here on imbalance in ERP amplitude, keeping the exact ERP shape constant (except for latency/amplitude noise that was comparable in both trial groups). However, systematic differences in ERP shape (in particular, in the onset or peak latency of specific components) are likely to also strongly affect phase opposition.

## References

Ai, L., & Ro, T. (2014). The phase of prestimulus alpha oscillations affects tactile perception. J Neurophysiol, 111(6), 1300–1307. doi:10.1152/jn.00125.2013

Arnal, L. H., Doelling, K. B., & Poeppel, D. (2015). Delta-Beta Coupled Oscillations Underlie Temporal Prediction Accuracy. Cereb Cortex, 25 (9), 3077–3085 doi:10.1093/cercor/bhu103

Auksztulewicz, R., & Blankenburg, F. (2013). Subjective Rating of Weak Tactile Stimuli Is Parametrically Encoded in Event-Related Potentials. Journal of Neuroscience, 33 (29), 11878–11887 doi:10.1523/Jneurosci.4243-12.2013

Barry, R. J., Rushby, J. A., Johnstone, S. J., Clarke, A. R., Croft, R. J., & Lawrence, C. A. (2004). Event-related potentials in the auditory oddball as a function of EEG alpha phase at stimulus onset. Clin Neurophysiol, 115 (11), 2593–2601 doi:10.1016/j.clinph.2004.06.004

Batterink, L. J., Creery, J. D., & Paller, K. A. (2016). Phase of Spontaneous Slow Oscillations during Sleep Influences Memory-Related Processing of Auditory Cues. Journal of Neuroscience, 36 (4), 1401–1409 doi:10.1523/Jneurosci.3175-15.2016

Baumgarten, T. J., Schnitzler, A., & Lange, J. (2015). Beta oscillations define discrete perceptual cycles in the somatosensory domain. Proc Natl Acad Sci U S A, 112 (39), 12187–12192 doi:10.1073/pnas. 1501438112

Benjamini, Y., & Hochberg, Y. (1995). Controlling the False Discovery Rate-a Practical and Powerful Approach to Multiple Testing. Journal of the Royal Statistical Society Series B-Methodological, 57 (1), 289–300

Berens, P. (2009). CircStat: A MATLAB Toolbox for Circular Statistics. Journal of Statistical Software, 31(10), 121.

Bompas, A., Sumner, P., Muthumumaraswamy, S. D., Singh, K. D., & Gilchrist, I. D. (2015). The contribution of pre-stimulus neural oscillatory activity to spontaneous response time variability. Neuroimage, 107, 34–45. doi:10.1016/j.neuroimage.2014.11.057

Bonnefond, M., & Jensen, O. (2012). Alpha oscillations serve to protect working memory maintenance against anticipated distracters. Curr Biol, 22 (20), 1969–1974

Busch, N. A., Dubois, J., & VanRullen, R. (2009). The phase of ongoing EEG oscillations predicts visual perception. J Neurosci, 29 (24), 7869–7876

Busch, N. A., & VanRullen, R. (2010). Spontaneous EEG oscillations reveal periodic sampling of visual attention. Proc Natl Acad Sci U S A, 107 (37), 16048–16053

Buschman, T. J., & Miller, E. K. (2009). Serial, covert shifts of attention during visual search are reflected by the frontal eye fields and correlated with population oscillations. Neuron, 63 (3), 386–396

Buzsaki, G. (2006). Rhythms of the Brain. New York: Oxford University Press.

Callaway, E. I., & Yeager, C. L. (1960). Relationship between reaction time and electroencephalographic alpha phase. Science, 132(1765-1766).

Chakravarthi, R., & VanRullen, R. (2012). Conscious updating is a rhythmic process. Proc Natl Acad Sci U S A, 109 (26), 10599–10604

Cravo, A. M., Santos, K. M., Reyes, M. B., Caetano, M. S., & Claessens, P. M. (2015). Visual Causality Judgments Correlate with the Phase of Alpha Oscillations. J Cogn Neurosci, 27 (10), 1887–1894 doi:10.1162/jocn_a_00832

Diederich, A., Schomburg, A., & van Vugt, M. (2014). Fronto-central theta oscillations are related to oscillations in saccadic response times (SRT): an EEG and behavioral data analysis. PLoS ONE, 9(11), e112974. doi:10.1371/journal.pone.0112974

Drewes, J., & VanRullen, R. (2011). This is the rhythm of your eyes: the phase of ongoing electroencephalogram oscillations modulates saccadic reaction time. J Neurosci, 31 (12), 4698–4708

Dugue, L., Marque, P., & VanRullen, R. (2011). The phase of ongoing oscillations mediates the causal relation between brain excitation and visual perception. J Neurosci, 31 (33), 11889–11893

Dugue, L., Marque, P., & VanRullen, R. (2015). Theta oscillations modulate attentional search performance periodically. J Cogn Neurosci, 27 (5), 945–958 doi:10.1162/jocn_a_00755

Dustman, R. E., & Beck, E. C. (1965). Phase of Alpha Brain Waves, Reaction Time and Visually Evoked Potentials. Electroencephalogr Clin Neurophysiol, 18, 433–440.

Fell, J., Dietl, T., Grunwald, T., Kurthen, M., Klaver, P., Trautner, P., … Fernandez, G. (2004). Neural bases of cognitive ERPs: More than phase reset. Journal of Cognitive Neuroscience, 16 (9), 1595–1604 doi:Doi 10.1162/0898929042568514

Fiebelkorn, I. C., Snyder, A. C., Mercier, M. R., Butler, J. S., Molholm, S., & Foxe, J. J. (2013). Cortical crossfrequency coupling predicts perceptual outcomes. Neuroimage, 69, 126–137. doi:10.1016/j.neuroimage.2012.11.021

Gruber, W. R., Zauner, A., Lechinger, J., Schabus, M., Kutil, R., & Klimesch, W. (2014). Alpha phase, temporal attention, and the generation of early event related potentials. Neuroimage, 103, 119–129. doi:10.1016/j.neuroimage.2014.08.055

Haig, A. R., & Gordon, E. (1998). EEG alpha phase at stimulus onset significantly affects the amplitude of the P3 ERP component. Int J Neurosci, 93(1-2), 101–115.

Hamm, J. P., Dyckman, K. A., McDowell, J. E., & Clementz, B. A. (2012). Pre-cue fronto-occipital alpha phase and distributed cortical oscillations predict failures of cognitive control. J Neurosci, 32 (20), 7034–7041

Han, B., & VanRullen, R. (2015). The rhythms of predictive coding: pre-stimulus oscillatory phase modulates the influence of shape perception on luminance judgments. (submitted).

Hanslmayr, S., Volberg, G., Wimber, M., Dalal, S. S., & Greenlee, M. W. (2013). Prestimulus oscillatory phase at 7 Hz gates cortical information flow and visual perception. Curr Biol, 23 (22), 2273–2278 doi:10.1016/j.cub.2013.09.020

Henry, M. J., Herrmann, B., & Obleser, J. (2014). Entrained neural oscillations in multiple frequency bands comodulate behavior. Proc Natl Acad Sci U S A, 111 (41), 14935–14940 doi:10.1073/pnas.1408741111

Inyutina, M., Sun, H. M., Wu, C. T., & VanRullen, R. (2015). Who wins the race for consciousness? Ask the phase of ongoing ~7Hz oscillations. J Vis, 15(12), 569. doi:10.1167/15.12.569

Jansen, B. H., & Brandt, M. E. (1991). The effect of the phase of prestimulus alpha activity on the averaged visual evoked response. Electroencephalogr Clin Neurophysiol, 80 (4), 241–250

Lachaux, J. P., Rodriguez, E., Martinerie, J., & Varela, F. J. (1999). Measuring phase synchrony in brain signals. Hum Brain Mapp, 8 (4), 194–208

Lakatos, P., Musacchia, G., O‘Connel, M. N., Falchier, A. Y., Javitt, D. C., & Schroeder, C. E. (2013). The spectrotemporal filter mechanism of auditory selective attention. Neuron, 77 (4), 750–761

Landau, A. N., Schreyer, H. M., van Pelt, S., & Fries, P. (2015). Distributed Attention Is Implemented through Theta-Rhythmic Gamma Modulation. Curr Biol, 25 (17), 2332–2337 doi:10.1016/j.cub.2015.07.048

Leszczynski, M., Fell, J., & Axmacher, N. (2015). Rhythmic Working Memory Activation in the Human Hippocampus. Cell Rep, 13 (6), 1272–1282 doi:10.1016/j.celrep.2015.09.081

Li, H., Zhang, L., Zhang, J. C., Wang, C. M., Yao, L., Wu, X., & Guo, X. J. (2015). Improving N1 classification by grouping EEG trials with phases of pre-stimulus EEG oscillations. Cognitive Neurodynamics, 9(2), 103-112. doi:10.1007/s11571-014-9317-x

Luck, S. J. (2014). An introduction to the event-related potential technique. Cambridge (MA): MIT Press.

Makeig, S., Westerfield, M., Jung, T. P., Enghoff, S., Townsend, J., Courchesne, E., & Sejnowski, T. J. (2002). Dynamic brain sources of visual evoked responses. Science, 295 (5555), 690–694

Manasseh, G., de Balthasar, C., Sanguinetti, B., Pomarico, E., Gisin, N., de Peralta, R. G., & Andino, S. L. G. (2013). Retinal and post-retinal contributions to the quantum efficiency of the human eye revealed by electrical neuroimaging. Frontiers in psychology, 4. doi:Artn 845, 10.3389/Fpsyg.2013.00845

Mathewson, K. E., Gratton, G., Fabiani, M., Beck, D. M., & Ro, T. (2009). To see or not to see: prestimulus alpha phase predicts visual awareness. J Neurosci, 29 (9), 2725–2732

Mazaheri, A., & Jensen, O. (2006). Posterior alpha activity is not phase-reset by visual stimuli. Proceedings of the National Academy of Sciences of the United States of America, 103 (8), 2948–2952 doi:10.1073/pnas.0505785103

McLelland, D., Lavergne, L., & VanRullen, R. (2014). The phase of ongoing EEG oscillations predicts the amplitude of peri-saccadic mislocalization. Paper presented at the Society for Neuroscience meeting, Washington DC.

Min, B. K., Busch, N. A., Debener, S., Kranczioch, C., Hanslmayr, S., Engel, A. K., & Herrmann, C. S. (2007). The best of both worlds: Phase-reset of human EEG alpha activity and additive power contribute to ERP generation. International Journal of Psychophysiology, 65 (1), 58–68 doi:10.1016/j.ijpsycho.2007.03.002

Moratti, S., Clementz, B. A., Gao, Y., Ortiz, T., & Keil, A. (2007). Neural mechanisms of evoked oscillations: Stability and interaction with transient events. Human Brain Mapping, 28 (12), 1318–1333 doi:10.1002/hbm. 20342

Myers, N. E., Stokes, M. G., Walther, L., & Nobre, A. C. (2014). Oscillatory brain state predicts variability in working memory. J Neurosci, 34 (23), 7735–7743 doi:10.1523/JNEUROSCI.4741-13.2014

Ng, B. S., Schroeder, T., & Kayser, C. (2012). A precluding but not ensuring role of entrained low-frequency oscillations for auditory perception. J Neurosci, 32 (35), 12268–12276

Nunn, C. M., & Osselton, J. W. (1974). The influence of the EEG alpha rhythm on the perception of visual stimuli. Psychophysiology, 11 (3), 294–303

Park, H. D., Correia, S., Ducorps, A., & Tallon-Baudry, C. (2014). Spontaneous fluctuations in neural responses to heartbeats predict visual detection. Nature neuroscience, 17(4), 612–U178. doi:10.1038/nn.3671

Rana, K. D., Vaina, L. M., & Hamalainen, M. S. (2013). A fast statistical significance test for baseline correction and comparative analysis in phase locking. Frontiers in Neuroinformatics, 7. doi:UNSP 3, 10.3389/fninf.2013.00003

Rice, D. M., & Hagstrom, E. C. (1989). Some evidence in support of a relationship between human auditory signal-detection performance and the phase of the alpha cycle. Percept Mot Skills, 69 (2), 451–457 doi:10.2466/pms.1989.69.2.451

Samaha, J., Bauer, P., Cimaroli, S., & Postle, B. R. (2015>). Top-down control of the phase of alpha-band oscillations as a mechanism for temporal prediction. Proc Natl Acad Sci U S A, 112 (27), 8439–8444 doi:10.1073/pnas.1503686112

Sauseng, P., Klimesch, W., Gruber, W. R., Hanslmayr, S., Frelinberger, R., & Doppelmayr, M. (2007). Are event-related potential components generated by phase resetting of brain oscillations? A critical discussion. Neuroscience, 146 (4), 1435–1444 doi:10.1016/j.neuroscience.2007.03.014

Scheeringa, R., Mazaheri, A., Bojak, I., Norris, D. G., & Kleinschmidt, A. (2011). Modulation of visually evoked cortical FMRI responses by phase of ongoing occipital alpha oscillations. J Neurosci, 31(10), 38133820.

Sherman, M. T., Kanai, R., Seth, A. K., & VanRullen, R. (2016). Rhythmic influence of top-down perceptual priors in the phase of pre-stimulus occipital alpha oscillations. J Cog Neuroscience, (in press).

Shou, G., & Ding, L. (2015). Detection of EEG Spatial-Spectral-Temporal Signatures of Errors: A Comparative Study of ICA-Based and Channel-Based Methods. Brain Topogr, 28 (1), 47–61 doi:10.1007/s10548-014-0397-z

Siegel, M., Warden, M. R., & Miller, E. K. (2009). Phase-dependent neuronal coding of objects in short-term memory. Proc Natl Acad Sci U S A, 106 (50), 21341–21346

Stouffer, S. A., Suchman, E. A., DeVinney, L. C., Star, S. A., & Williams, R. M. J. (1949). Studies in Social Psychology in World War II: The American Soldier. Vol. 1, Adjustment During Army Life. Princeton: Princeton University Press.

Strauss, A., Henry, M. J., Scharinger, M., & Obleser, J. (2015). Alpha phase determines successful lexical decision in noise. J Neurosci, 35 (7), 3256–3262 doi:10.1523/JNEUROSCI.3357-14.2015

Stroud, J. M. (1956). The fine structure of psychological time. In H. Quastler (Ed.), Information theory in Psychology (pp. 174-205). Chicago, Ill: Free Press.

Ten Oever, S., van Atteveldt, N., & Sack, A. T. (2015). Increased Stimulus Expectancy Triggers Low-frequency Phase Reset during Restricted Vigilance. J Cogn Neurosci, 27 (9), 1811–1822 doi:10.1162/jocn_a_00820

van Diepen, R. M., Cohen, M. X., Denys, D., & Mazaheri, A. (2015). Attention and temporal expectations modulate power, not phase, of ongoing alpha oscillations. J Cogn Neurosci, 27 (8), 1573–1586 doi:10.1162/jocn_a_00803

van Erp, J. B., Philippi, T. G., de Winkel, K. N., & Werkhoven, P. (2014). Pre-and post-stimulus EEG patterns associated with the touch-induced illusory flash. Neurosci Lett, 562, 79–84. doi:10.1016/j.neulet.2014.01.010

VanRullen, R., Busch, N. A., Drewes, J., & Dubois, J. (2011). Ongoing EEG phase as a trial-by-trial predictor of perceptual and attentional variability. Frontiers in Perception Science, 2 (60), 1–9

VanRullen, R., & Koch, C. (2003). Is perception discrete or continuous? Trends Cogn Sci, 7 (5), 207–213

Varela, F. J., Toro, A., John, E. R., & Schwartz, E. L. (1981). Perceptual framing and cortical alpha rhythm. Neuropsychologia, 19 (5), 675–686

Vinck, M., van Wingerden, M., Womelsdorf, T., Fries, P., & Pennartz, C. M. (2010). The pairwise phase consistency: a bias-free measure of rhythmic neuronal synchronization. Neuroimage, 51 (1), 112–122

Voloh, B., Valiante, T. A., Everling, S., & Womelsdorf, T. (2015). Theta-gamma coordination between anterior cingulate and prefrontal cortex indexes correct attention shifts. Proc Natl Acad Sci U S A, 112 (27), 8457–8462 doi:10.1073/pnas.1500438112

Wyart, V., de Gardelle, V., Scholl, J., & Summerfield, C. (2012). Rhythmic fluctuations in evidence accumulation during decision making in the human brain. Neuron, 76 (4), 847–858 doi:10.1016/j.neuron.2012.09.015

Zar, J. H. (1999). Biostatistical analysis: Prentice Hall.

Zoefel, B., & Heil, P. (2013). Detection of Near-Threshold Sounds is Independent of EEG Phase in Common Frequency Bands. Frontiers in psychology, 4, 262.

